# Regional Signals Matter: Rethinking Global Pooling for mRNA Foundation Models

**DOI:** 10.64898/2026.11.12.766253

**Authors:** Saiyang Na, Zhijie Qi, Deevanshu Goyal, Junzhou Huang, Gene Yeo, Wenhao Jin

## Abstract

RNA foundation models with diverse architectures have shown promise for learning representations of messenger RNA (mRNA) sequences. However, existing benchmarks commonly reduce token-level embeddings to transcript-level representations through global mean pooling before fitting a downstream probe. This aggregation does not explicitly preserve the identities of biologically distinct mRNA regions. Consequently, signals associated with short regulatory regions may be diluted, confounding assessments of representation quality with the information preserved by the readout. We introduce an architecture-agnostic, region-aware readout that separately averages embeddings within the 5^′^ untranslated region (UTR), coding sequence, and 3^′^ UTR and concatenates the resulting representations for linear probing. We reproduce mRNABench evaluations for 20 frozen models from 18 families, with a median absolute score difference of 0.0009 across 380 model–dataset–target evaluations published for 19 models, and construct a region-aware benchmark using the same models, tasks, and splits. Relative to global mean pooling, region-aware pooling increases Pearson correlation, averaged across the 20 models, by 11–25% on translation and stability tasks, with no statistically significant decreases detected for any model on the regression tasks. On classification tasks, relative changes in the area under the precision–recall curve range from −3% to +11%. Boundary randomization and displacement generally reduce performance, while dimensionality-matched and sequence-composition controls indicate that the gains are not explained solely by increased feature dimensionality, region length, or guanine–cytosine (GC) content. Learned attention pooling approaches region-aware performance only when supplied with one query per region. In silico perturbations reveal stronger sensitivity to upstream AUGs (uAUGs) and their Kozak context, while analyses of endogenous transcripts show approximately threefold greater recovery of length- and GC-adjusted uAUG- associated differences than global-mean probes. Both readouts capture codon optimality. Model rankings averaged over the regression tasks remain largely stable, while within a task models move by several places. Our region-aware benchmark therefore provides a biologically structured and complementary evaluation of mRNA foundation models, revealing regulatory information that is less accessible to linear probes after global mean pooling. The code is available at https://github.com/Sanford-Labs/rnaregionpooling.

## 1 Introduction

RNA foundation models learn transferable representations from large collections of unlabeled nucleotide sequences, providing a common framework for predicting translation, RNA stability, localization, and other molecular properties from sequence. Models applied to messenger RNA (mRNA) span diverse architectures and pretraining strategies, including bidirectional Transformers trained on coding and non-coding RNA, DNA language models transferred to transcripts, long- context state-space and Hyena-based models, and models pre- trained specifically on mature mRNAs with transcript annota- tions [1, 2, 3, 4, 5, 6, 7]. Their utility is commonly evaluated by freezing the encoder, extracting token-level embeddings, and training a lightweight probe to predict experimentally measured properties such as translation efficiency (TE), mean ribosome load (MRL), RNA half-life (HL), or subcellular lo- calization [8]. For transcript-level probing, token embeddings are commonly aggregated into a single vector by global mean pooling.

An mRNA comprises biologically distinct regions with different regulatory functions. The 5^′^ untranslated region (5^′^ UTR) influences translation initiation through upstream AUG codons (uAUGs) and upstream open reading frames (uORFs), while sequence context around the translation start site further modulates initiation [9, 10, 11]. The coding sequence (CDS) contains codon-usage patterns associated with elongation and mRNA decay [12], and the 3^′^ UTR contains AU-rich elements (AREs), microRNA (miRNA) target sites, and localization signals [13, 14]. Global mean pooling gives longer regions greater direct weight in the pooled representation. Across the evaluated mRNABench datasets, median 5^′^ UTR lengths range from 160 to 210 nucleotides, and these regions account for approximately 5–10% of transcript length. Although contextual token embeddings may encode positional and regional information, global averaging does not explicitly preserve region identity. A downstream probe must therefore recover localized regulatory signals from an aggregated representation. Benchmark performance consequently reflects both the information encoded by the model and its accessibility under the chosen aggregation and probing scheme.

Related approaches include region-specific pretraining [15, 16, 17], separate regional encoders [18], annotation- aware modeling [6, 7] and learned aggregation [19]. Learned pooling fits its aggregation parameters to downstream labels, so its performance can reflect both the pretrained representations and the additional capacity of the aggregation module; fitted on one model, attention pooling comes within 3% of region-aware pooling only when given one query per region (Table 1). These approaches address related but distinct questions. Here, we ask whether preserving annotated region identity during aggregation improves access to regulatory information in existing frozen representations, without modifying the encoder or learning aggregation parameters.

**Table 1:** Learned attention pooling on RiNALMo giga token embeddings for TE human (ten splits, early stopping on validation), compared with the ridge probes; the highest test r is bold. Share of the pooling weight: the part of the weight on each region, mean over the test transcripts and the ten splits; under the nucleotide mean, the regions’ shares of the nucleotides. NA, not applicable (the readout pools each region or third on its own); SD, standard deviation.

| **Readout** | **Test r** | **SD over splits** | **Share of the pooling weight** |  |  |
|---|---|---|---|---|---|
|  |  |  | **5′ UTR** | **CDS** | **3′ UTR** |
| Attention pooling, no region information | 0.6521 | 0.0117 | 0.045 | 0.660 | 0.294 |
| Attention pooling, region-type embedding on the keys | 0.6577 | 0.0125 | 0.078 | 0.622 | 0.300 |
| Attention pooling, one query per region | 0.7121 | 0.0081 | NA | NA | NA |
| Ridge, nucleotide mean | 0.6672 | 0.0120 | 0.075 | 0.498 | 0.427 |
| Ridge, equal thirds | 0.6731 | 0.0132 | NA | NA | NA |
| Ridge, region-aware | **0.7316** | 0.0098 | NA | NA | NA |

We address this question with region-aware pooling, which averages the embeddings of a frozen model within the 5^′^ UTR, CDS, and 3^′^ UTR and concatenates the three regional means (Figure 1). As the reference, we reproduce mRNABench’s readout and linear probes for 20 models from 18 families on its ten splits, and we evaluate region-aware pooling on the same models, tasks, and splits.

**Figure 1:**
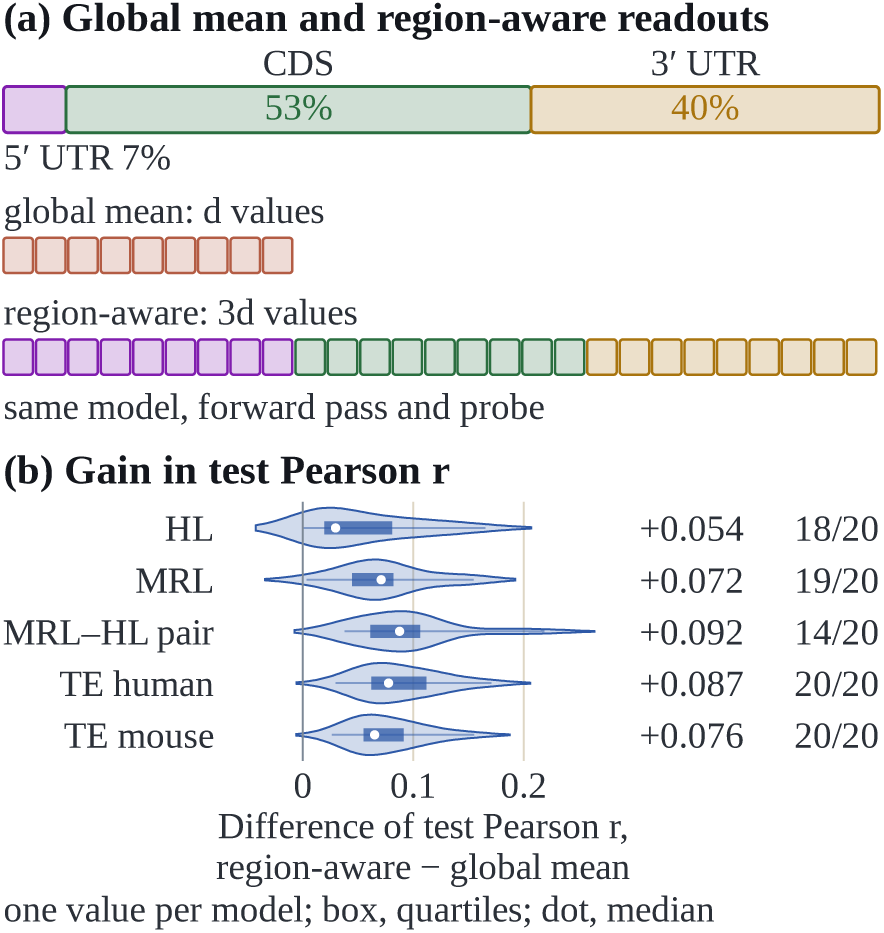
Region-aware and global mean pooling. (a) A transcript’s three regions, their median share of its nucleotides, and the two readouts of one frozen model’s token embeddings. (b) Difference of test Pearson r, region-aware minus global mean pooling, mean over the 20 models; the fractions count the models in which it is positive at Wilcoxon p < 0.05 over the ten splits.

Concatenating regional representations increases feature dimensionality and, for a standard linear probe, the number of fitted coefficients.

Our contributions are as follows.

1. **A parameter-free regional aggregation scheme for frozen mRNA representations.** Region-aware pooling reuses existing encoder outputs, introduces no trainable pooling parameters, and leaves the pretrained model unchanged. It applies to architectures whose output embeddings can be aligned to annotated nucleotide regions and uses region coordinates already available in the benchmark datasets.
2. **Controlled evaluation of the sources of performance gains.** We test whether improvements depend on annotated biological boundaries rather than increased feature dimensionality, regional length and composi- tion, or encoder-layer choice. The evaluation includes equal-width, randomized, and shifted segmentations; dimensionality-matched readouts; length and guanine–cytosine (GC) content features; independently encoded regions; alternative encoder layers; and learned attention pooling.
3. **Biological analyses of regulatory signals accessible under different readouts.** We combine in silico pertur- bations of uAUGs, synonymous codons, AREs, and miRNA target sites with analyses of natural transcripts and regional decomposition of linear predictions.
4. **Implications for mRNA foundation-model benchmarking.** Predictive scores and task-specific model rankings depend on the readout. Reporting region-aware pooling alongside global mean pooling provides a complemen- tary evaluation that helps distinguish encoder performance from readout-dependent accessibility of regulatory information.

## 2 Related Work

### 2.1 Foundation models for RNA and mRNA

RNA language models trained with masked language modeling on non-coding RNA (ncRNA) databases include RNABERT, RNA-FM, RNA-MSM, RNAErnie, ERNIE-RNA and RiNALMo [20, 1, 21, 22, 23, 2]. SpliceBERT is trained on pre-mRNA sequences from 72 vertebrates [24], and AIDO.RNA scales masked language modeling on ncRNA to 1.6B parameters [25]. Region-specific models restrict pretraining to one part of the transcript: UTR-LM to 5^′^ UTRs, 3UTRBERT to 3^′^ UTRs and CodonBERT to coding sequences [15, 16, 17]. DNA language models, among them DNABERT-2, DNABERT-S, the Nucleotide Transformer, HyenaDNA, Evo and Evo 2, can be applied to transcripts as nucleotide strings [26, 27, 3, 4, 28, 5]. Models trained on mature mRNA take the regions as input or combine region encoders: Orthrus’s six-track input marks coding positions and splice sites, Helix-mRNA inserts a mark token before every codon [6, 7], and mRNA-LM combines three region encoders [18]. Apart from mRNA-LM these models are read out with a transcript-level mean or a single token. We change only the readout and use the models as released; Table S1 lists the 20 we evaluate.

### 2.2 Benchmarks and readouts

Benchmarks for nucleotide models collect common tasks so that architectures can be compared on equal terms [29, 30]; BEND and mRNABench read the models out frozen and fit a small head or a linear probe on the embeddings [31, 8]. mRNABench collects mature-mRNA tasks — massively parallel reporter assays (MPRAs) on 5^′^ UTRs, endogenous ribosome load and translation efficiency, half-life, localization, RNA binding and variant effects — and probes each model’s global mean with ridge or logistic regression on ten splits [8]; its published scores are the reference we reproduce. Readouts have received less attention than models. Mean pooling is the standard reduction for protein language models; light attention, locality-aware bag-of-mer pooling and optimal-transport pooling are the learned alternatives [19, 32, 33], a comparison of compression methods found mean pooling hard to beat [34], and covariance pooling has been proposed for a genomic model [35]. On mRNABench tasks, RIBOSPAN encodes inputs longer than a model’s context as separate chunks averaged by length and attributes the gap between its 1K- and 10K-context models to this chunking [36]. These studies treat the transcript, or the protein, as one unit. We ask whether the unit should be the region.

### 2.3 Region-aware modeling of mRNA

Supervised predictors of mRNA properties use the regions as separate inputs or features: Optimus 5-Prime and Framepool read ribosome load off the 5^′^ UTR alone [37, 38], Saluki predicts half-life from the full transcript with coding-frame and splice-site tracks [39], and RiboNN attributes about 22%, 73% and 5% of its translation-efficiency predictions to the 5^′^ UTR, the CDS and the 3^′^ UTR [40]. mRNA-LM is the closest construction to ours: it encodes each region with its own model and trains on the concatenated region means [18], so each region is encoded without the others and the encoders and the head are trained, where region-aware pooling keeps one frozen model and trains only the probe. We know of no study that pools a frozen full-length model by region and compares that across model families.

### 2.4 Regulatory elements and in silico perturbation

The elements we perturb have well-characterized effects. An upstream AUG or uORF reduces translation of the main open reading frame (ORF), more so in a strong Kozak context, whereas an in-frame AUG without an intervening stop extends the protein [9, 10, 37]. Codon optimality, the association of a codon with longer mRNA half-life, is a major determinant of mRNA stability [12]. AREs, of which UUAUUUAUU is the core nonamer, shorten half-life [41, 42, 43], as do miRNA target sites in 3^′^ UTRs, whose efficacy follows 8mer > 7mer-m8 > 7mer-A1 > 6mer [14, 44]. In silico mutagenesis is a standard way to test whether a trained predictor responds to such elements [45, 37, 39]. We apply it to linear probes on frozen embeddings, with matched controls, and complement it with groupings of natural transcripts by the elements they already carry, so that a probe’s response can be compared with the measured effect in the same dataset.

## 3 Methods

### 3.1 Datasets and data splits

We use the datasets and the ten data splits of mRNABench release 1.3 [8] (Table 2). The regression columns are mean ribosome load (MRL) in four 5^′^ UTR MPRA libraries (designed, mCherry, varying length, and eGFP with uridine, pseudouridine or N1-methylpseudouridine; MRL MPRA) [37], MRL of endogenous mRNA isoforms in RCC4 cells re-expressing the von Hippel–Lindau protein, resolved by transcription start site (MRL) [46], half-life (HL) in human and mouse [39], ribosome load and in-cell half-life of 203 designed mRNAs (MRL–HL pair) [47], and translation efficiency (TE) in human and mouse (TE human, TE mouse) [40]. The classification columns are variant effect prediction (VEP) [48], RNA-binding protein (RBP) binding measured by enhanced crosslinking and immunoprecipitation (eCLIP) [49, 50], mRNA localization from long-read and short-read sequencing (Loc-LR, Loc- SR) [51, 52], protein localization (Prot-Loc) [53] and Gene Ontology (GO) terms [54]. Our Loc-LR labels come from this release and are not compared with the published column. Each dataset is split ten times 70/15/15 with mRNABench’s seeds, by gene homology group for the endogenous datasets and at random for the MPRA libraries, the MRL–HL pair and VEP; for the mouse half-life dataset we follow mRNABench’s choice of homology map. A column’s score is the mean over its datasets, with each dataset’s targets averaged first, computed per split; we report the mean over the ten splits.

**Table 2:** Evaluation columns, their source datasets in mRNABench and the measured quantities. A column is the mean over its datasets, each dataset’s targets averaged first. Score: Pearson r for the regression columns and the area under the precision–recall curve (AUPRC) for the classification columns.

| **Column** | **Benchmark table** | **Transcripts** | **Measured property** | **Score** |
|---|---|---|---|---|
| HL | rnahl-human, rnahl-mouse | 26,662 | half-life | Pearson r |
| MRL | mrl-sugimoto | 11,693 | ribosome load | Pearson r |
| MRL MPRA | mrl-sample-designed, mrl-sample-egfp, mrl-sample-mcherry, mrl-sample-varying | 716,812 | ribosome load | Pearson r |
| MRL–HL pair | mrl-hl-lbkwk | 203 | ribosome load, half-life | Pearson r |
| TE human | translation-efficiency-human | 10,281 | translation efficiency | Pearson r |
| TE mouse | translation-efficiency-mouse | 10,853 | translation efficiency | Pearson r |
| VEP | vep-traitgym-complex, vep-traitgym-mendelian |  | variant effect | AUPRC |
| eCLIP | eclip-binding-hepg2, eclip-binding-k562 |  | protein binding | AUPRC |
| Loc-LR, Loc-SR | rna-lifecycle-ietswaart, rna-loc-fazal |  | localization | AUPRC |
| Prot-Loc | prot-loc |  | protein localization | AUPRC |
| GO | go-bp, go-cc, go-mf |  | gene ontology | AUPRC |

### 3.2 Models and embeddings

We evaluate 20 frozen models from 18 families (Table S1): RNA language models (RNABERT, RNA-FM, RNA- MSM, RNAErnie, ERNIE-RNA, RiNALMo giga, SpliceBERT, AIDO.RNA 1.6B), region-specific models (UTR-LM, 3UTRBERT), models trained on mature mRNA (Orthrus large 6-track and base 4-track, Helix-mRNA and HelixNano- mRNA, of which the 6-track input and Helix-mRNA’s codon marks carry annotation) and DNA language models (DNABERT-2, DNABERT-S, the multi-species Nucleotide Transformer 2.5B (NT 2.5B), HyenaDNA, Evo 1, Evo 2 7B in its 8K-context base form) [20, 1, 21, 22, 23, 2, 24, 25, 15, 16, 6, 7, 55, 26, 27, 3, 4, 28, 5]. Each transcript (5^′^ UTR, CDS and 3^′^ UTR) is embedded with the last layer of the model, using mRNABench’s tokenization, special tokens, chunking of long inputs and numerical precision for that model. We refer to the benchmark’s readout as the global mean: mRNABench’s mean of the token vectors, computed as each of its model wrappers computes it.

### 3.3 Region-aware pooling

From the same forward pass, each nucleotide is assigned the vector of the token that covers it, and special tokens are dropped. Let ℎ*_i_* be the vector at nucleotide *i*, and *R*_5_, *R_C_* and *R*_3_ the positions of the three regions in the dataset’s annotation. The region-aware readout is the concatenation

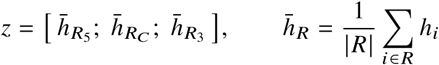

a vector three times the model’s hidden size; a region that a transcript lacks contributes zeros. Figure 2 compares it with the global mean. The readout adds no parameters and needs only the region coordinates the benchmark datasets carry.

**Figure 2:**
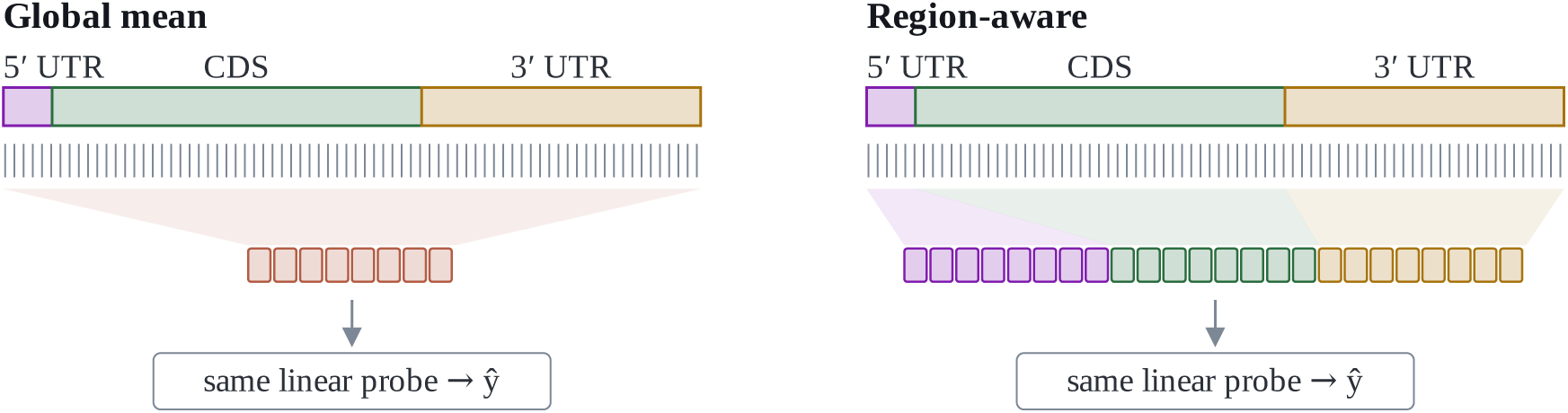
Global and region-aware pooling of a frozen model’s token embeddings. Ticks denote the per-nucleotide embeddings of the frozen model. The global mean pooling averages all of them into d values and weights each region by its length: the 5^′^ UTR, 5–10% of the nucleotides, contributes about a twentieth of the vector. Region-aware pooling averages within each region and concatenates the three means into 3d values, each region with its own d coordinates. The same linear probe is fitted on either vector.

### 3.4 Linear probing and statistical testing

For regression columns we use ridge regression with an unpenalized intercept, choosing the penalty by exact leave- one-out error on the training rows; classification columns use mRNABench’s logistic regression (C = 1) and are scored by the area under the precision–recall curve (AUPRC). The global mean is probed exactly as in mRNABench. Our readouts use the same estimators, with the ridge penalty searched over 10^−6^ to 10^6^. All comparisons are paired by split. For each model and column, the ten split-wise differences are tested with a two-sided Wilcoxon signed-rank test, and the model counts as a win or a loss at p < 0.05 and as a tie otherwise.

### 3.5 Control readouts

Equal thirds cuts each transcript into three parts of equal length, ignoring the annotation, and concatenates their means, which gives a vector of the same width as the region readout. The nucleotide mean averages the per-nucleotide vectors over the whole transcript. In the principal component analysis (PCA) control, both readouts are reduced to their top d principal components on the training rows, d being the model’s hidden size. The balanced mean averages the three region means, giving each region equal weight in a vector of the model’s hidden size. The length and GC features (log lengths and GC fractions of the three regions) are appended to a readout. Random boundaries place the two boundaries at the region proportions of another transcript drawn at random (five draws, averaged); shifted boundaries move both boundaries by 10, 30, 100 or 300 nt toward either end. Per-region encoding feeds each region to the model on its own instead of the whole transcript at once (15 models).

### 3.6 Decomposition of predictions by region

For the region probe with weights w = (*w*_5_, *w_C_*, *w*_3_) and test rows x = (*x*_5_, *x_C_*, *x*_3_), the prediction is a sum of three segment contributions *c_k_* = (*x_k_* − *μ_k_*) · *w_k_*, with *μ_k_* the mean of *x_k_* over the training rows. For each segment we report the variance of its contribution over the test rows as a share of the prediction’s variance; the remaining share is the covariance between segments.

### 3.7 Sensitivity to regulatory elements

In silico mutagenesis. We draw transcripts from the default test split of a dataset and edit them by length-preserving substitution [45, 37]. An AUG is inserted into the 5^′^ UTR as an N-terminal extension (in frame with the main ORF, with no stop before it), an overlapping uORF (oORF: out of frame, with no stop before the annotated start codon) or a uORF (closed by an in-frame stop), each in a strong Kozak context (a purine at −3 and a G at +4) or a weak one (neither). Each edit has a control in which a non-AUG codon is substituted at the same position, and in transcripts with a single uAUG we change that AUG to AUC (TE human, MRL, eGFP MPRA). On HL human, a quarter, half or all of the codons are recoded to their most optimal, least optimal or a random synonym according to the codon stability coefficient (CSC) of the training genes [12], and four copies of the ARE nonamer UUAUUUAUU or of a let-7 8mer site are inserted into the 3^′^ UTR at least 15 nt past the stop codon, each beside a shuffled control [42, 14, 44]. Probes are fitted on the default train split, and the response to an edit is the mean change of the prediction in units of the label’s standard deviation (SD). As the measured reference, transcripts whose 5^′^ UTR carries exactly one uAUG are grouped by kind and Kozak context and compared with the uAUG-free transcripts of the same dataset; a probe’s response is summarized by its slope on these six measured class effects.

Natural groupings. Without edits, the test rows of each split are grouped by an element they carry: the uAUG classes; out-of-frame and in-frame uAUGs; the top and bottom quartiles of mean codon optimality (CSC) in the CDS and of the codon adaptation index (CAI) [56]; three or more AUUUA motifs in the 3^′^ UTR [41]; and let-7 or miR-16 sites (TargetScan 8mer, 7mer-m8 and 7mer-A1) [44]. For each grouping we compare the group difference in the measured label with that in each probe’s held-out predictions, both raw and adjusted for the log length and GC of the three regions, and report the share of the measured difference that the probe reproduces. A probe that captured every aspect of the label equally would reproduce about r^2^ of any group difference, r being its test correlation; we show this share as a reference.

## 4 Results

### 4.1 Reproduction of the mRNABench baselines

For the 19 models that mRNABench reports (HelixNano-mRNA is not among them), our global-mean scores match the published ones with a median absolute difference of 0.0009, or 0.2% of the published score, over 380 model– dataset–target cells, and 325 cells differ by at most 0.01. The 42 cells that differ by more than 0.02 are concentrated in RNAErnie (17), whose released weights have been replaced since the benchmark was run, UTR-LM (8), Helix-mRNA (7) and Evo 1 (5); for Evo 1, mRNABench rounds the mean to bfloat16, and its MPRA and pair columns respond to small numerical differences. All comparisons below are made against this reproduction, on the same splits.

### 4.2 Region-aware pooling improves translation and stability prediction

Table 3 summarizes the twelve columns. The region readout scores higher than the global mean on every regression column, and the three counts give the models in which the difference is positive or negative at Wilcoxon p < 0.05 and those in which it is not significant: TE human +0.087 (20/0/0), TE mouse +0.076 (20/0/0), MRL +0.072 (19/0/1), HL +0.054 (18/0/2), MRL–HL pair +0.092 (14/0/6) and MRL MPRA +0.059 (20/0/0), or 11–25% of the global mean. No model is significantly worse on any regression column (Table S2); Table S3 gives every model under every readout. On the classification columns the changes range from −0.008 to +0.028 AUPRC (−3% to +11%).

**Table 3:** Mean over 20 models (regression: test Pearson r; classification: AUPRC). The global mean is mRNABench’s readout and probe as published; the other three are ours. The last two columns compare the region-aware readout with the global mean: the difference (region-aware minus global mean) and the number of models whose ten-split difference is positive / negative at Wilcoxon p < 0.05 / not significant. The best readout in each row is bold.

| **Column** | **Metric** | **Global mean** | **Nucleotide mean** | **Equal thirds** | **Region-aware** | **Region-aware vs global mean** |  |
|---|---|---|---|---|---|---|---|
|  |  |  |  |  |  | **difference** | **better / worse / tied** |
| ***Local*** |  |  |  |  |  |  |  |
| VEP | AUPRC | 0.2485 | 0.2471 | 0.2702 | **0.2763** | +0.0278 | 13 / 2 / 5 |
| MRL MPRA | Pearson r | 0.5209 | 0.5402 | 0.5759 | **0.5802** | +0.0593 | 20 / 0 / 0 |
| eCLIP | AUPRC | 0.3508 | 0.3535 | **0.3710** | 0.3684 | +0.0176 | 14 / 5 / 1 |
| ***Global (RNA)*** |  |  |  |  |  |  |  |
| Loc-LR | AUPRC | 0.5211 | 0.5230 | **0.5297** | 0.5278 | +0.0067 | 11 / 6 / 3 |
| Loc-SR | AUPRC | 0.6784 | 0.6826 | 0.6816 | **0.6858** | +0.0075 | 10 / 6 / 4 |
| HL | Pearson r | 0.4649 | 0.4831 | 0.5019 | **0.5191** | +0.0541 | 18 / 0 / 2 |
| MRL | Pearson r | 0.2841 | 0.2962 | 0.3273 | **0.3564** | +0.0723 | 19 / 0 / 1 |
| MRL–HL pair | Pearson r | 0.4541 | 0.4770 | 0.4918 | **0.5457** | +0.0916 | 14 / 0 / 6 |
| TE human | Pearson r | 0.6152 | 0.6303 | 0.6538 | **0.7018** | +0.0866 | 20 / 0 / 0 |
| TE mouse | Pearson r | 0.5763 | 0.5925 | 0.6124 | **0.6521** | +0.0758 | 20 / 0 / 0 |
| ***Global (protein)*** |  |  |  |  |  |  |  |
| Prot-Loc | AUPRC | 0.3237 | **0.3273** | 0.3111 | 0.3152 | −0.0085 | 9 / 9 / 2 |
| GO | AUPRC | 0.3279 | 0.3316 | 0.3235 | **0.3341** | +0.0062 | 13 / 7 / 0 |

### 4.3 The gain depends on the annotated region boundaries

Table 4 compares our readouts with each other on the regression columns. At equal width, the region readout beats equal thirds by 0.017–0.054, and it keeps a lead of 0.018–0.054 when both are reduced to the model’s hidden size. Random boundaries remove the gain: HL, MRL, TE human and TE mouse drop by 0.053–0.085, and all 20 models drop on each of these columns. Shifting both boundaries 100 nt toward the 5^′^ end moves the last 100 nt before the start codon into the CDS segment and reduces MRL by 0.041 in all 20 models; the shift toward the 3^′^ end leaves the 5^′^ UTR intact and reduces it by 0.008. With region lengths and GC appended to both readouts, the region readout stays 0.021–0.055 above the nucleotide mean given the same six features. Averaging the three region means into one vector (the balanced mean) does not reproduce the gain, which therefore depends on keeping the regions apart. Encoding each region on its own, without the context of the other two, raises the region readout further: by 0.007 on MRL, 0.016 and 0.020 on TE human and mouse and 0.053 on the 203 designed mRNAs of the MRL–HL pair. Of the 15 models the rise is significant in 11 (TE human), 10 (TE mouse) and 9 (MRL, pair); HL rises 0.005, significantly in 5 models and in reverse in 4. Pooled from one whole-transcript pass, the region readout keeps 80–91% of what per-region encoding gains over the global mean on HL, MRL and TE. The region boundaries matter when the model encodes the transcript as well as when its output is pooled. The gain is not specific to the last layer: taken from the middle layer of ERNIE-RNA, RNAErnie, RNA-FM, SpliceBERT and RiNALMo giga, the region readout stays above the global mean on the five regression columns other than MRL MPRA in all five models (0.023–0.143, median 0.090), and by a wider margin than at the last layer in 20 of the 25 model–column pairs.

**Table 4:** Control readouts on the regression columns: mean difference over models, with the same three counts as in Table 3. Per-region encoding was run for 15 models.

| Control | HL | MRL | MRL–HL pair | TE human | TE mouse |
|---|---|---|---|---|---|
| ***Difference from equal thirds*** |  |  |  |  |  |
| Region-aware | +0.0171 17 / 1 / 2 | +0.0291 16 / 0 / 4 | +0.0539 14 / 0 / 6 | +0.0480 19 / 1 / 0 | +0.0396 18 / 2 / 0 |
| Region-aware, both reduced to d by PCA | +0.0181 19 / 1 / 0 | +0.0343 17 / 0 / 3 | +0.0539 14 / 0 / 6 | +0.0497 19 / 1 / 0 | +0.0429 19 / 1 / 0 |
| ***Difference from the nucleotide mean*** |  |  |  |  |  |
| Balanced region mean | −0.0197 1 / 17 / 2 | +0.0149 13 / 3 / 4 | +0.0441 12 / 0 / 8 | +0.0054 13 / 5 / 2 | −0.0071 9 / 7 / 4 |
| Region-aware, both with length and GC | +0.0206 16 / 0 / 4 | +0.0553 18 / 0 / 2 | +0.0500 9 / 1 / 10 | +0.0342 20 / 0 / 0 | +0.0309 19 / 0 / 1 |
| ***Difference from annotated boundaries*** |  |  |  |  |  |
| Random boundaries | −0.0534 0 / 20 / 0 | −0.0846 0 / 20 / 0 | −0.0358 0 / 11 / 9 | −0.0807 0 / 20 / 0 | −0.0746 0 / 20 / 0 |
| Shifted 100 nt toward the 5′ end | −0.0069 1 / 16 / 3 | −0.0413 0 / 20 / 0 | −0.0727 0 / 16 / 4 | −0.0210 1 / 19 / 0 | −0.0246 0 / 19 / 1 |
| Shifted 100 nt toward the 3′ end | −0.0047 0 / 12 / 8 | −0.0082 1 / 10 / 9 | +0.0063 2 / 4 / 14 | −0.0096 1 / 17 / 2 | −0.0080 2 / 13 / 5 |
| ***Difference from per-region encoding*** |  |  |  |  |  |
| Whole-transcript encoding | −0.0051 4 / 5 / 6 | −0.0073 4 / 9 / 2 | −0.0534 1 / 9 / 5 | −0.0164 3 / 11 / 1 | −0.0200 3 / 10 / 2 |

### 4.4 Regional contributions to the predictions

The decomposition of the region probe follows the regulatory roles of the regions (Figure 3). Ribosome load draws on the 5^′^ UTR (median share 0.56 over the 20 models, 0.48 on the transcripts that have all three regions), half-life on the CDS (0.54) and the 3^′^ UTR (0.36), and translation efficiency on the CDS (0.51 human, 0.53 mouse) and the 5^′^ UTR (0.27 and 0.29). In the global mean the 5^′^ UTR has a weight of 5–10%, its share of the nucleotides; in the region probe, its segment carries more than half of the ribosome-load prediction.

**Figure 3:**
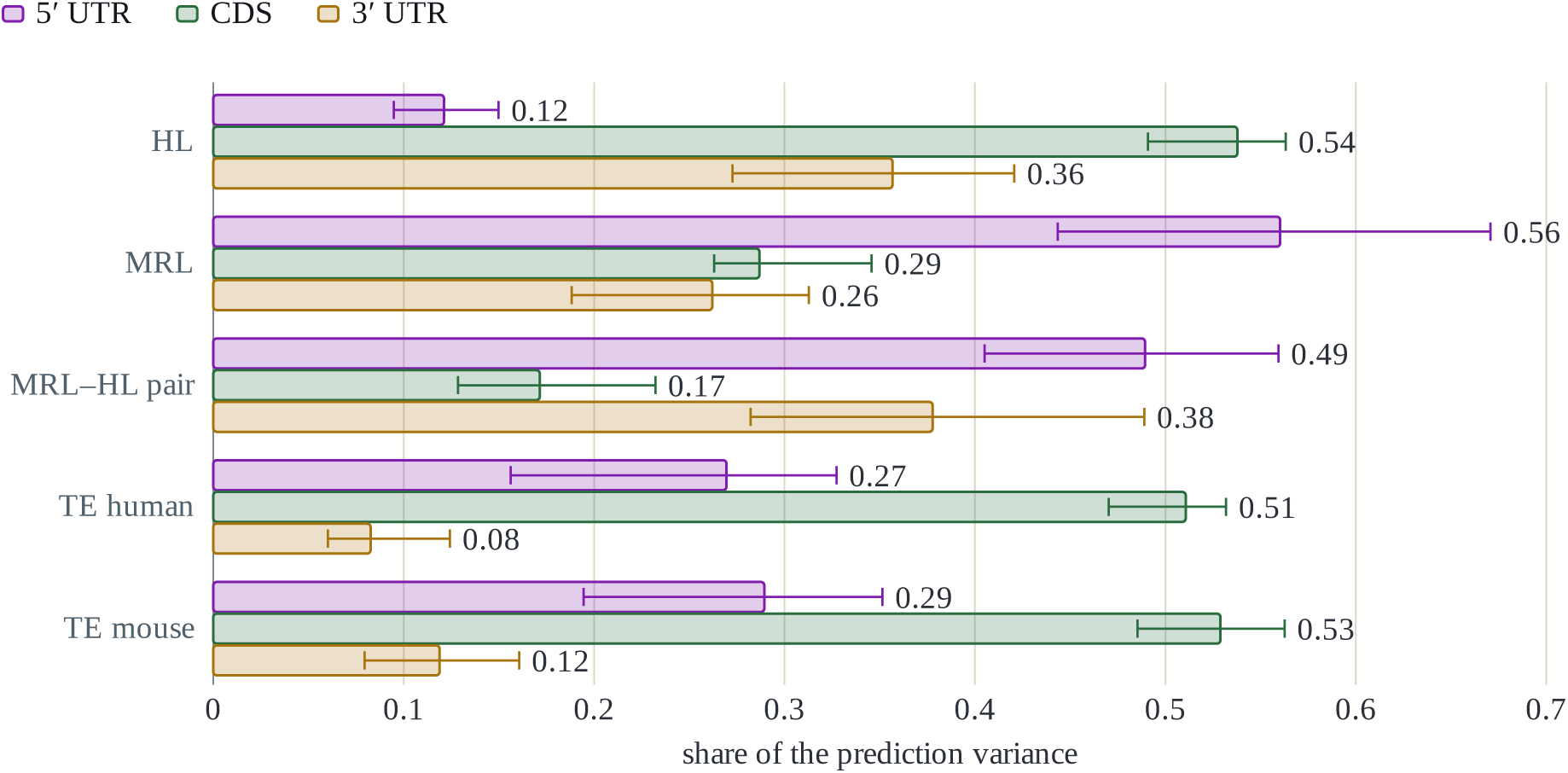
Share of the region-aware probe’s prediction variance carried by each segment: median over the 20 models (bar) and interquartile range (whiskers); the remainder is covariance between segments.

### 4.5 Region-aware probes capture the effects of upstream AUGs

An informative readout should allow the probe to respond to elements with a known effect, in the measured direction and with the measured size, and leave it unchanged under matched neutral edits. We first group natural transcripts by their uAUGs (Figure 4, Table S4). Transcripts carrying a single uAUG that opens an oORF or a uORF, or several uAUGs, have a measured TE and MRL 0.20–0.91 SD below those of uAUG-free transcripts; weak-context uORFs on TE mouse and MRL are the exception and sit near zero. After adjustment for region length and GC, the held-out predictions of the global mean reproduce 7–22% of these differences and those of the region readout 24–63%, and the region readout is closer to the measurement in 18–20 of the 20 models for each class. Accuracy alone does not explain this. On the TE datasets the global mean stays well below its r^2^ (0.34–0.39), while the region readout approaches its own (0.43–0.50); on MRL the region readout exceeds its r^2^ (0.13) by 1.8- to 2.8-fold. In the four MPRA libraries, where only the 5^′^ UTR varies, out-of-frame uAUGs lower MRL by 1.4–1.6 SD after adjustment. The global mean reproduces 40–44% of this effect and the region readout 51–55%, and the region readout is closer in all 20 models on every library. In-frame uAUGs have a milder measured effect (−0.33 to −0.42 SD), which both readouts reproduce at or above its measured size (global mean 90–105%, region readout 107–129%; Figure S1). Both readouts thus recover the stronger repression associated with out-of-frame than in-frame uAUGs, consistent with previously reported frame-dependent effects on ribosome loading [37].

**Figure 4:**
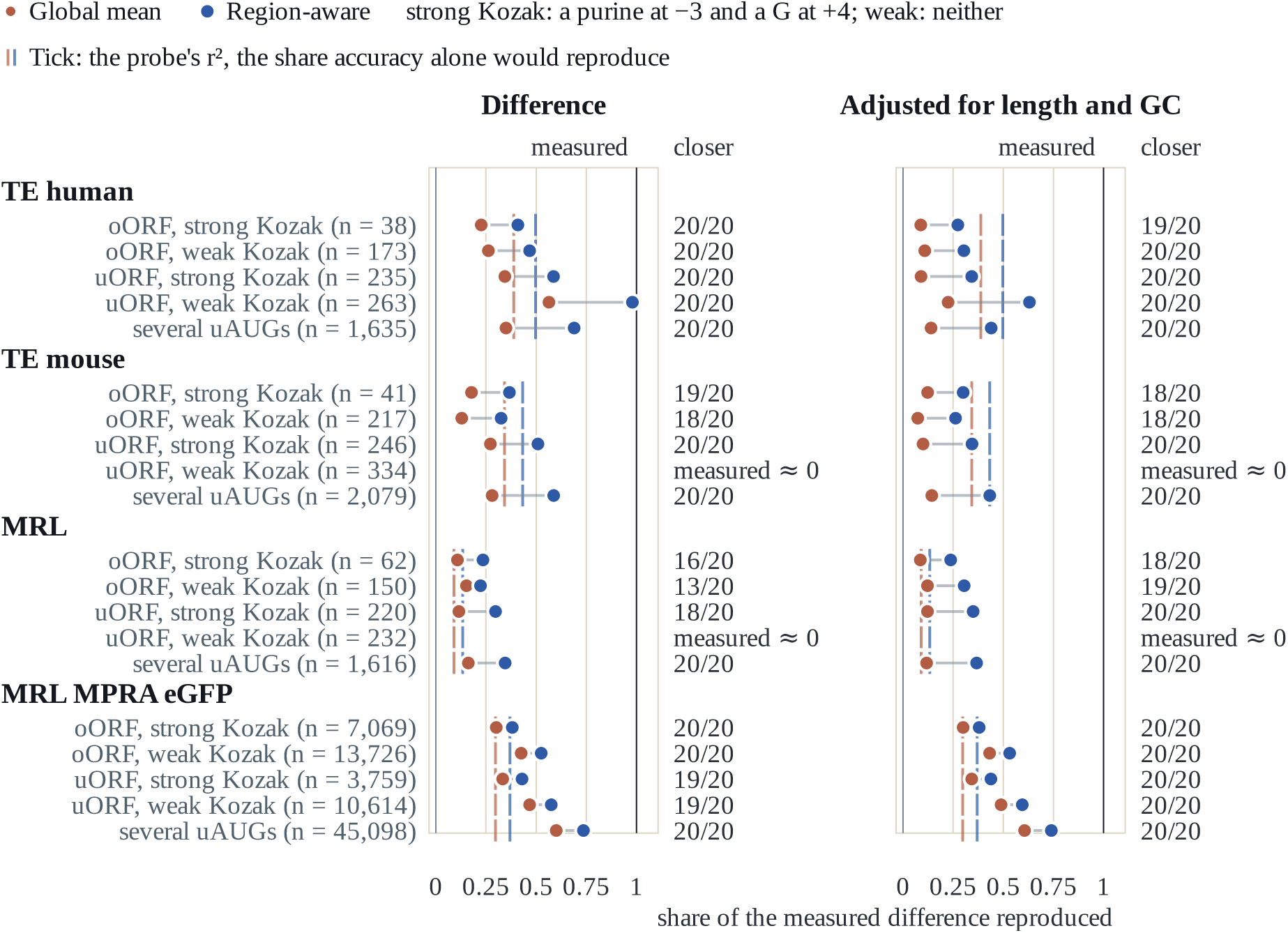
Measured and predicted effects of upstream AUGs in natural transcripts. Each row is a group of test transcripts whose 5^′^ UTR carries a single uAUG of one class, or several uAUGs (n in parentheses), against the uAUG- free transcripts of the same dataset. A marker is the share of the measured group difference that a probe’s held-out predictions reproduce — the predicted difference over the measured one, both in units of the label’s standard deviation, averaged over the ten splits and then over the 20 models — so 1 (black line) is the full measured effect, 0 no effect and above 1 an overestimate. Red dots are the global mean, blue dots the region-aware readout. Left, the raw differences; right, the same differences adjusted for the log length and GC content of the three regions. A short tick marks a probe’s r^2^, its mean squared test Pearson r: the share of any group difference it would reproduce if it caught every aspect of the label equally, so a marker right of its tick reproduces more of that class than the probe’s accuracy alone accounts for. The right-hand column counts the models in which the region-aware readout lands closer to the measurement. Classes measured within 0.15 SD of the uAUG-free transcripts carry no markers.

Inserting a uAUG into transcripts (Figure 5, Table S5) yields consistent results in 20 models. On the endogenous datasets the global-mean prediction moves by at most 0.10 SD and reproduces at most 15% of the measured effect (median about 0% on TE human and 1% on MRL). Region-aware predictions decrease by up to 0.29 SD in a strong Kozak context and by less in a weak context. These changes correspond to a median of 9% and 15% of the observed group differences on TE human and MRL, respectively, reaching up to 32% and 49%. The 5^′^ UTR component accounts for a median of 92% and 102% of the total predicted change on the two datasets, respectively; values above 100% reflect opposing contributions from the other regions. Prediction changes remain within 0.03 SD of zero under control edits and increase by up to 0.15 SD upon uAUG ablation. The region readout reproduces more of the measured effect than the global mean in 18 of the 20 models on both datasets. On the eGFP MPRA, where only the 5^′^ UTR varies, both readouts follow the measured order of the six classes (median *ρ* 0.91 for the global mean and 0.94 for the region readout), and the region readout reproduces more of the effect in all 20 models (median slope 0.44 against 0.31).

**Figure 5:**
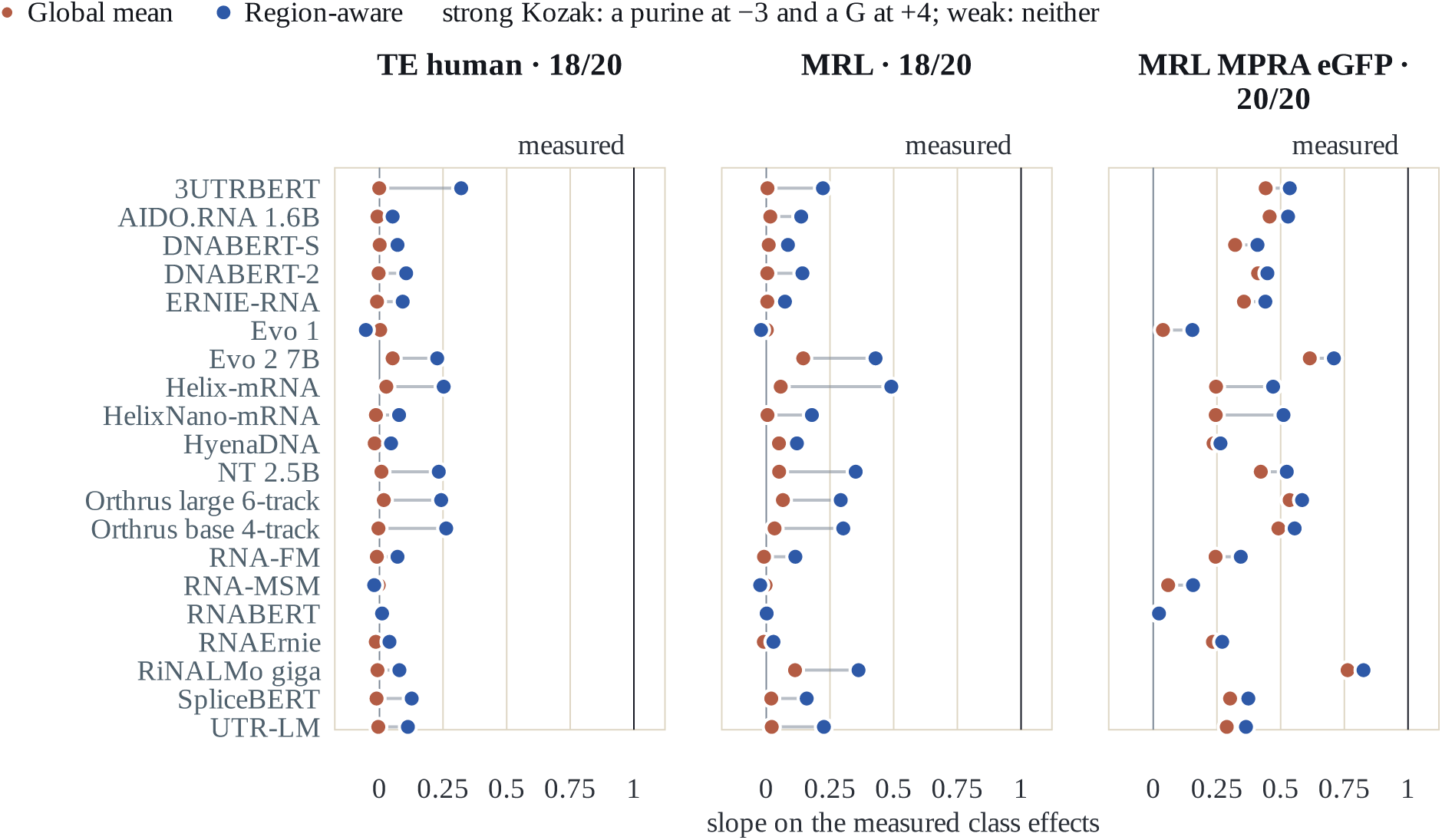
In silico insertion of upstream AUGs. A uAUG is substituted into the 5^′^ UTR of each test transcript in each of the six classes of Figure 4, and the change in each probe’s prediction is recorded. A marker is the slope of that mean predicted change, in units of the label’s standard deviation, regressed on the six measured class effects of the same dataset, so that 1 (black line) means the probe moves exactly as much as the class effects measured on natural transcripts and 0 that it does not move. Red dots are the global mean, blue dots the region-aware readout; one row per model, one panel per dataset. Each panel title gives the dataset and the number of models whose region-aware slope is the closer of the two to 1.

### 4.6 Regulatory elements in the CDS and the 3^′^ UTR

The CDS and the 3^′^ UTR make up most of a transcript, so the global mean does not dilute them. Codon optimality has strong measured effects (Figure S1), +0.24 to +0.93 SD between the top and bottom quartiles after adjustment. The global mean reproduces 59–74% of this difference and the region readout 70–99%, both far above their r^2^; the region readout is closer in 3 to 11 of the 20 models and the global mean in at most one.

Synonymous recoding changes the CDS alone. Recoding a quarter, half or all of the codons to their most or least optimal synonyms (Table S7 gives the quarter and the whole) moves the global-mean probe in the expected direction in 19 of the 20 models and the region probe in 15 to 17, and the global mean about twice as far (with a quarter of the codons recoded, which keeps the transcripts within the natural range of codon optimality: +0.23 against +0.10 SD between optimal and least optimal synonyms; Figure S2). Length weighting also accounts for this result, here in favor of the global mean. The CDS holds about half of the nucleotides of an HL human transcript (median 0.53), and in natural transcripts its codon optimality correlates with the composition of the 3^′^ UTR (r = 0.59 between mean CSC and 3^′^ UTR GC). In the global mean the two regions share one vector, so the composition signal its probe learns from natural transcripts spans both, and a recoded CDS moves that signal as a whole. The region probe keeps the CDS and the 3^′^ UTR in separate segments, which carry 0.54 and 0.36 of its half-life prediction (Figure 3), and a recoded CDS moves only the first. The two readouts are thus complementary: the global mean is the more sensitive to composition spread over the longest region, the region readout to elements in the shortest.

For AUUUA motifs and miRNA target sites (Table S8), both probes follow the part of the difference that region length and GC explain. After adjustment, the measured AUUUA effect halves to −0.23 to −0.26 SD and the miRNA effects shrink to −0.01 to −0.10; the global mean reproduces 50–68% of the adjusted AUUUA difference and the region readout 32–61%, neither readout reproduces more than a sixth of the adjusted miRNA differences, and four inserted AREs or let-7 sites move the predictions no more than shuffled controls do (differences of at most 0.10 SD).

### 4.7 Effect of the readout on model rankings

The gain is consistent across models, whereas the model ranking is not. Table 5 gives each model’s scores under the region readout. Within a column the two readouts order the 20 models with Kendall *τ* between 0.31 and 0.86, and on none of the twelve columns is the order unchanged: 8 to 19 models take a different place (Figure S3, Table S9). 3UTRBERT, which is pretrained on 3^′^ UTRs, rises on 11 of the 12 columns and by as much as seven places; RNA- FM, UTR-LM, Helix-mRNA and HelixNano-mRNA rise on seven to ten columns, while NT 2.5B, RiNALMo giga, AIDO.RNA 1.6B and DNABERT-S fall on eight or more, by up to thirteen places. On the regression columns no model scores significantly lower under the region readout (Table S2), so the global mean underestimates the models that move up. Averaged over the five regression columns other than MRL MPRA, the two orders are close (*τ* 0.88, 12 of the 20 models keep their place); an aggregate score thus masks the per-task reordering.

**Table 5:**
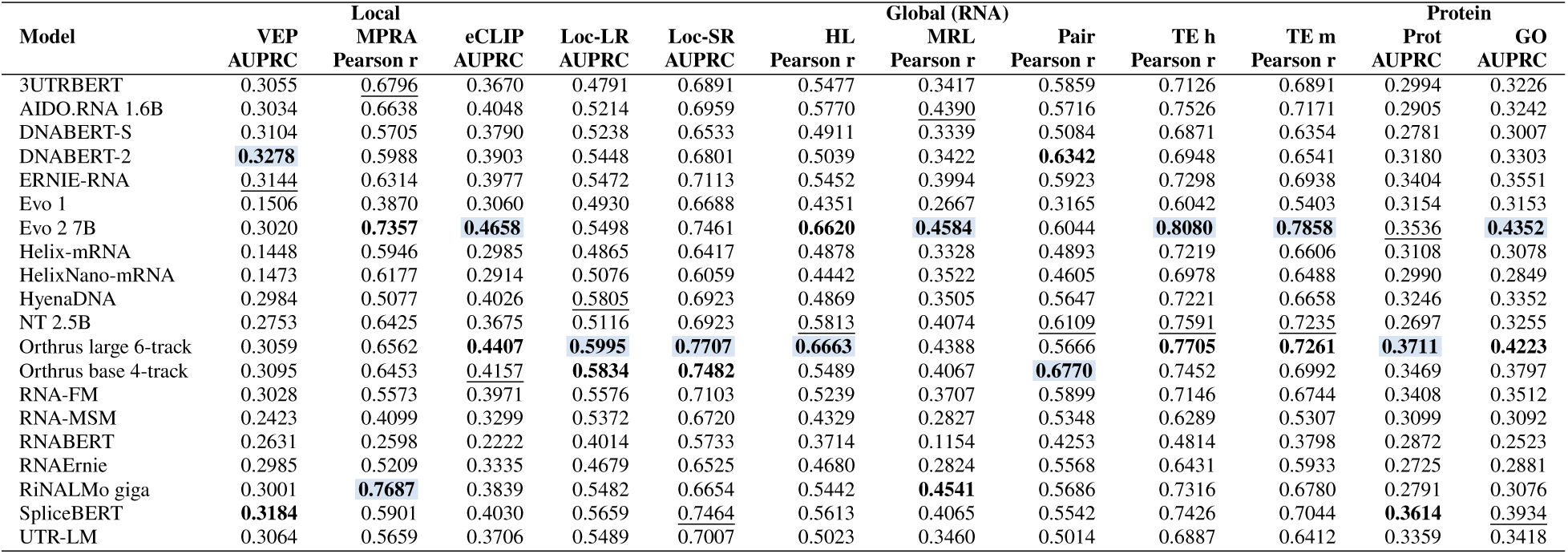
Region-aware readout per model and column, mean over the ten splits (regression: test Pearson r; classification: AUPRC). Top three per column: **shaded and bold** , **bold**, underlined; Table S3 gives every readout. MPRA, MRL MPRA; Pair, the MRL–HL pair; TE h and TE m, TE human and TE mouse; Prot, Prot-Loc.

|  | Local |  |  | Global (RNA) |  |  |  |  |  |  | Protein |  |
|---|---|---|---|---|---|---|---|---|---|---|---|---|
| **Model** | **VEP** | **MPRA** | **eCLIP** | **Loc-LR** | **Loc-SR** | **HL** | **MRL** | **Pair** | **TE h** | **TE m** | **Prot** | **GO** |
|  | **AUPRC** | **Pearson r** | **AUPRC** | **AUPRC** | **AUPRC** | **Pearson r** | **Pearson r** | **Pearson r** | **Pearson r** | **Pearson r** | **AUPRC** | **AUPRC** |
| 3UTRBERT | 0.3055 | 0.6796 | 0.3670 | 0.4791 | 0.6891 | 0.5477 | 0.3417 | 0.5859 | 0.7126 | 0.6891 | 0.2994 | 0.3226 |
| AIDO.RNA 1.6B | 0.3034 | 0.6638 | 0.4048 | 0.5214 | 0.6959 | 0.5770 | 0.4390 | 0.5716 | 0.7526 | 0.7171 | 0.2905 | 0.3242 |
| DNABERT-S | 0.3104 | 0.5705 | 0.3790 | 0.5238 | 0.6533 | 0.4911 | 0.3339 | 0.5084 | 0.6871 | 0.6354 | 0.2781 | 0.3007 |
| DNABERT-2 | **0.3278** | 0.5988 | 0.3903 | 0.5448 | 0.6801 | 0.5039 | 0.3422 | **0.6342** | 0.6948 | 0.6541 | 0.3180 | 0.3303 |
| ERNIE-RNA | 0.3144 | 0.6314 | 0.3977 | 0.5472 | 0.7113 | 0.5452 | 0.3994 | 0.5923 | 0.7298 | 0.6938 | 0.3404 | 0.3551 |
| Evo 1 | 0.1506 | 0.3870 | 0.3060 | 0.4930 | 0.6688 | 0.4351 | 0.2667 | 0.3165 | 0.6042 | 0.5403 | 0.3154 | 0.3153 |
| Evo 2 7B | 0.3020 | **0.7357** | **0.4658** | 0.5498 | 0.7461 | **0.6620** | **0.4584** | 0.6044 | **0.8080** | **0.7858** | 0.3536 | **0.4352** |
| Helix-mRNA | 0.1448 | 0.5946 | 0.2985 | 0.4865 | 0.6417 | 0.4878 | 0.3328 | 0.4893 | 0.7219 | 0.6606 | 0.3108 | 0.3078 |
| HelixNano-mRNA | 0.1473 | 0.6177 | 0.2914 | 0.5076 | 0.6059 | 0.4442 | 0.3522 | 0.4605 | 0.6978 | 0.6488 | 0.2990 | 0.2849 |
| HyenaDNA | 0.2984 | 0.5077 | 0.4026 | 0.5805 | 0.6923 | 0.4869 | 0.3505 | 0.5647 | 0.7221 | 0.6658 | 0.3246 | 0.3352 |
| NT 2.5B | 0.2753 | 0.6425 | 0.3675 | 0.5116 | 0.6923 | 0.5813 | 0.4074 | 0.6109 | 0.7591 | 0.7235 | 0.2697 | 0.3255 |
| Orthrus large 6-track | 0.3059 | 0.6562 | **0.4407** | **0.5995** | **0.7707** | **0.6663** | 0.4388 | 0.5666 | **0.7705** | **0.7261** | **0.3711** | **0.4223** |
| Orthrus base 4-track | 0.3095 | 0.6453 | 0.4157 | **0.5834** | **0.7482** | 0.5489 | 0.4067 | **0.6770** | 0.7452 | 0.6992 | 0.3469 | 0.3797 |
| RNA-FM | 0.3028 | 0.5573 | 0.3971 | 0.5576 | 0.7103 | 0.5239 | 0.3707 | 0.5899 | 0.7146 | 0.6744 | 0.3408 | 0.3512 |
| RNA-MSM | 0.2423 | 0.4099 | 0.3299 | 0.5372 | 0.6720 | 0.4329 | 0.2827 | 0.5348 | 0.6289 | 0.5307 | 0.3099 | 0.3092 |
| RNABERT | 0.2631 | 0.2598 | 0.2222 | 0.4014 | 0.5733 | 0.3714 | 0.1154 | 0.4253 | 0.4814 | 0.3798 | 0.2872 | 0.2523 |
| RNAErnie | 0.2985 | 0.5209 | 0.3335 | 0.4679 | 0.6525 | 0.4680 | 0.2824 | 0.5568 | 0.6431 | 0.5933 | 0.2725 | 0.2881 |
| RiNALMo giga | 0.3001 | **0.7687** | 0.3839 | 0.5482 | 0.6654 | 0.5442 | **0.4541** | 0.5686 | 0.7316 | 0.6780 | 0.2791 | 0.3076 |
| SpliceBERT | **0.3184** | 0.5901 | 0.4030 | 0.5659 | 0.7464 | 0.5613 | 0.4065 | 0.5542 | 0.7426 | 0.7044 | **0.3614** | 0.3934 |
| UTR-LM | 0.3064 | 0.5659 | 0.3706 | 0.5489 | 0.7007 | 0.5023 | 0.3460 | 0.5014 | 0.6887 | 0.6412 | 0.3359 | 0.3418 |

## 5 Discussion

Global mean pooling weights regions by length, potentially diluting signals from shorter regions, whose relative lengths vary across transcripts. Region-aware pooling preserves region identity and improves access to regional signals. Gains concentrate on translation and stability tasks, depend on annotated boundaries, and include greater sensitivity to uAUGs. However, global pooling retains distributed signals such as codon usage and responds more strongly to synonymous CDS recoding. The readouts therefore complement each other rather than uniformly favoring regional pooling.

For benchmarking, these findings have two implications. First, global-mean scores can understate the accessibility of regional regulatory information. Absolute scores improve while aggregate rankings remain largely stable, although task-specific rankings shift. The uAUG analyses demonstrate this principle and provide a framework for evaluating other elements across the 5^′^ UTR, CDS, and 3^′^ UTR; benefits are not inherently restricted to the 5^′^ UTR. Second, annotated region coordinates make this complementary evaluation inexpensive: region-aware pooling requires only an additional pass over precomputed embeddings. Reporting both readouts can therefore help distinguish encoder performance from readout-dependent accessibility of regulatory information.

## 6 Conclusion

Global pooling dilutes the regulatory signals of short mRNA regions. Pooling within the 5^′^ UTR, the CDS and the 3^′^ UTR, with the same frozen model, forward pass and linear probe, improves translation and stability prediction, with no model significantly worse, depends on the annotated boundaries, and makes the probe sensitive to upstream AUGs.

## Supplementary material

The tables and figures below list the models and hold the per-model numbers behind the results, the uAUG classes and insertions, the HL edits, every element grouping and the model rankings.

**Table S1:** The 20 frozen models, their model class (an RNA, DNA or mRNA model, one pretrained on a single region, or an mRNA model that takes annotation as input) and the hidden size d of one token vector. The region-aware readout is 3d wide.

| **Model** | **Kind** | **Hidden size d** | **Model** | **Kind** | **Hidden size d** |
|---|---|---|---|---|---|
| 3UTRBERT | RNA, one region | 768 | NT 2.5B | DNA | 2,560 |
| AIDO.RNA 1.6B | RNA | 2,048 | Orthrus base 4-track | mRNA | 256 |
| DNABERT-2 | DNA | 768 | Orthrus large 6-track | mRNA, annotated | 512 |
| DNABERT-S | DNA | 768 | RNA-FM | RNA | 640 |
| ERNIE-RNA | RNA | 768 | RNA-MSM | RNA | 768 |
| Evo 1 | DNA | 4,096 | RNABERT | RNA | 120 |
| Evo 2 7B | DNA | 8,192 | RNAErnie | RNA | 768 |
| Helix-mRNA | mRNA, annotated | 256 | RiNALMo giga | RNA | 1,280 |
| HelixNano-mRNA | mRNA | 256 | SpliceBERT | RNA | 512 |
| HyenaDNA | DNA | 256 | UTR-LM | RNA, one region | 128 |

**Table S2:**
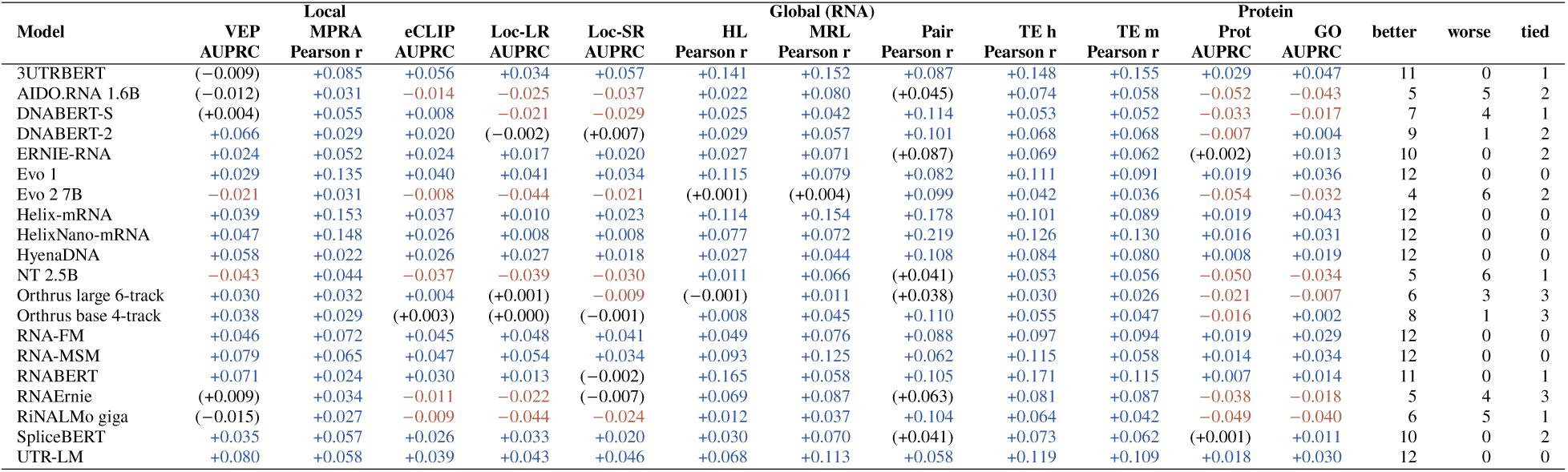
Region-aware readout minus global mean, per model and column (ten-split mean). At Wilcoxon p < 0.05, blue: region-aware higher, red: global mean higher; in parentheses: not significant. The last three columns count the columns in which each model is better, worse or neither at Wilcoxon p < 0.05. Short column names as in Table 5.

| Model | VEP | MPRA | eCLIP | Loc-LR | Loc-SR | HL | MRL | Pair | TE h | TE m | Prot | GO | better | worse | tied |
|---|---|---|---|---|---|---|---|---|---|---|---|---|---|---|---|
|  | Local |  |  |  |  | Global (RNA) |  |  |  |  | Protein |  |  |  |  |
| Model | VEP | MPRA | eCLIP | Loc-LR | Loc-SR | HL | MRL | Pair | TE h | TE m | Prot | GO | better | worse | tied |
|  | AUPRC | Pearson r | AUPRC | AUPRC | AUPRC | Pearson r | Pearson r | Pearson r | Pearson r | Pearson r | AUPRC | AUPRC |  |  |  |
| 3UTRBERT | (−0.009) | +0.085 | +0.056 | +0.034 | +0.057 | +0.141 | +0.152 | +0.087 | +0.148 | +0.155 | +0.029 | +0.047 | 11 | 0 | 1 |
| AIDO.RNA 1.6B | (−0.012) | +0.031 | −0.014 | −0.025 | −0.037 | +0.022 | +0.080 | (+0.045) | +0.074 | +0.058 | −0.052 | −0.043 | 5 | 5 | 2 |
| DNABERT-S | (+0.004) | +0.055 | +0.008 | −0.021 | −0.029 | +0.025 | +0.042 | +0.114 | +0.053 | +0.052 | −0.033 | −0.017 | 7 | 4 | 1 |
| DNABERT-2 | +0.066 | +0.029 | +0.020 | (−0.002) | (+0.007) | +0.029 | +0.057 | +0.101 | +0.068 | +0.068 | −0.007 | +0.004 | 9 | 1 | 2 |
| ERNIE-RNA | +0.024 | +0.052 | +0.024 | +0.017 | +0.020 | +0.027 | +0.071 | (+0.087) | +0.069 | +0.062 | (+0.002) | +0.013 | 10 | 0 | 2 |
| Evo 1 | +0.029 | +0.135 | +0.040 | +0.041 | +0.034 | +0.115 | +0.079 | +0.082 | +0.111 | +0.091 | +0.019 | +0.036 | 12 | 0 | 0 |
| Evo 2 7B | −0.021 | +0.031 | −0.008 | −0.044 | −0.021 | (+0.001) | (+0.004) | +0.099 | +0.042 | +0.036 | −0.054 | −0.032 | 4 | 6 | 2 |
| Helix-mRNA | +0.039 | +0.153 | +0.037 | +0.010 | +0.023 | +0.114 | +0.154 | +0.178 | +0.101 | +0.089 | +0.019 | +0.043 | 12 | 0 | 0 |
| HelixNano-mRNA | +0.047 | +0.148 | +0.026 | +0.008 | +0.008 | +0.077 | +0.072 | +0.219 | +0.126 | +0.130 | +0.016 | +0.031 | 12 | 0 | 0 |
| HyenaDNA | +0.058 | +0.022 | +0.026 | +0.027 | +0.018 | +0.027 | +0.044 | +0.108 | +0.084 | +0.080 | +0.008 | +0.019 | 12 | 0 | 0 |
| NT 2.5B | −0.043 | +0.044 | −0.037 | −0.039 | −0.030 | +0.011 | +0.066 | (+0.041) | +0.053 | +0.056 | −0.050 | −0.034 | 5 | 6 | 1 |
| Orthrus large 6-track | +0.030 | +0.032 | +0.004 | (+0.001) | −0.009 | (−0.001) | +0.011 | (+0.038) | +0.030 | +0.026 | −0.021 | −0.007 | 6 | 3 | 3 |
| Orthrus base 4-track | +0.038 | +0.029 | (+0.003) | (+0.000) | (−0.001) | +0.008 | +0.045 | +0.110 | +0.055 | +0.047 | −0.016 | +0.002 | 8 | 1 | 3 |
| RNA-FM | +0.046 | +0.072 | +0.045 | +0.048 | +0.041 | +0.049 | +0.076 | +0.088 | +0.097 | +0.094 | +0.019 | +0.029 | 12 | 0 | 0 |
| RNA-MSM | +0.079 | +0.065 | +0.047 | +0.054 | +0.034 | +0.093 | +0.125 | +0.062 | +0.115 | +0.058 | +0.014 | +0.034 | 12 | 0 | 0 |
| RNABERT | +0.071 | +0.024 | +0.030 | +0.013 | (−0.002) | +0.165 | +0.058 | +0.105 | +0.171 | +0.115 | +0.007 | +0.014 | 11 | 0 | 1 |
| RNAErnie | (+0.009) | +0.034 | −0.011 | −0.022 | (−0.007) | +0.069 | +0.087 | (+0.063) | +0.081 | +0.087 | −0.038 | −0.018 | 5 | 4 | 3 |
| RiNALMo giga | (−0.015) | +0.027 | −0.009 | −0.044 | −0.024 | +0.012 | +0.037 | +0.104 | +0.064 | +0.042 | −0.049 | −0.040 | 6 | 5 | 1 |
| SpliceBERT | +0.035 | +0.057 | +0.026 | +0.033 | +0.020 | +0.030 | +0.070 | (+0.041) | +0.073 | +0.062 | (+0.001) | +0.011 | 10 | 0 | 2 |
| UTR-LM | +0.080 | +0.058 | +0.039 | +0.043 | +0.046 | +0.068 | +0.113 | +0.058 | +0.119 | +0.109 | +0.018 | +0.030 | 12 | 0 | 0 |

**Table S3:**
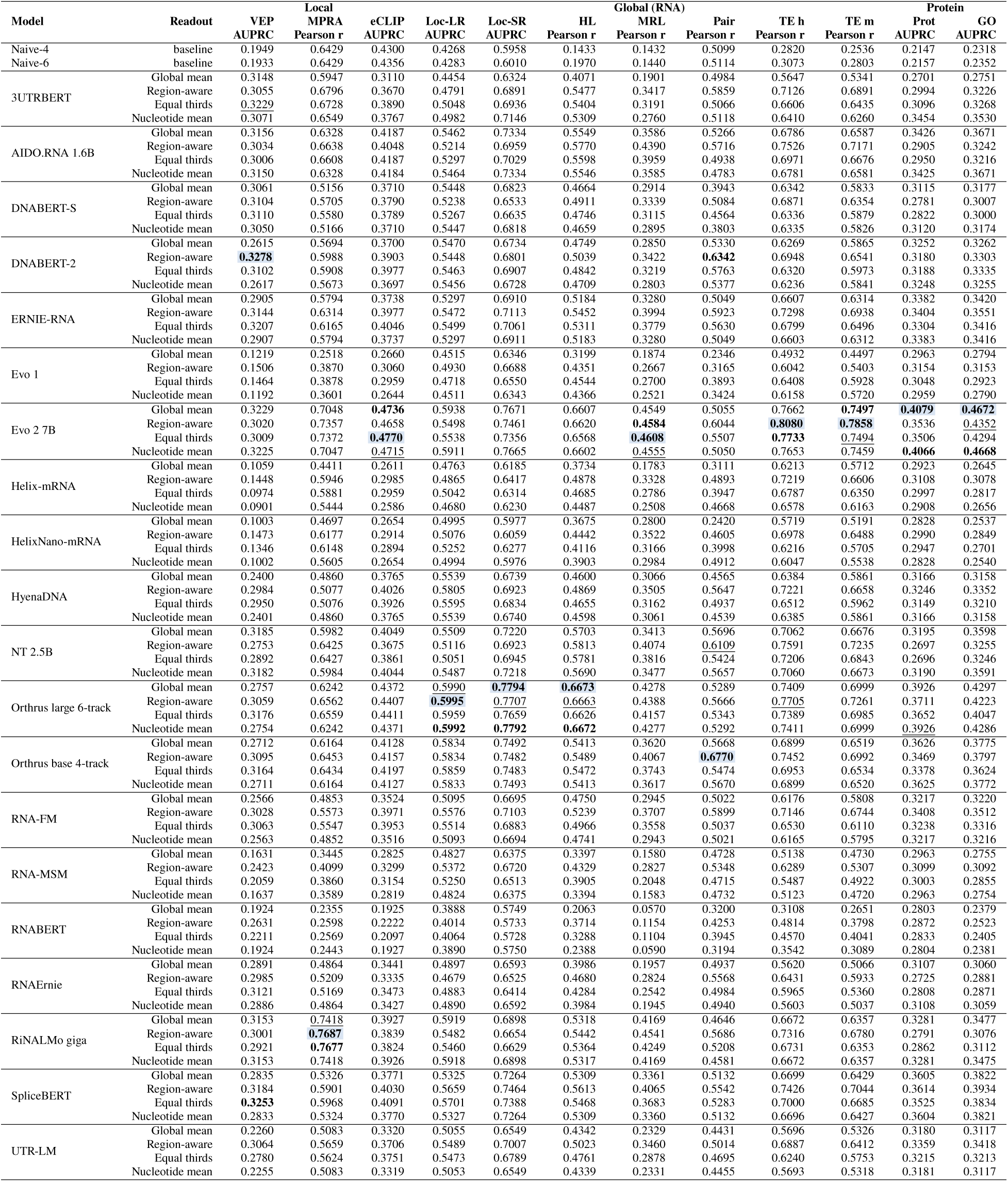
Ten-split means for every model and column, one row per readout. In each column the best of all 80 model– readout rows is **shaded and bold** , the second **bold** and the third underlined. Naive-4 and Naive-6 are mRNABench’s baselines on four and six sequence features. Short column names as in Table 5.

| **Model** | **Readout** | **Local** |  |  | **Global (RNA)** |  |  |  |  |  |  | **Protein** |  |
|---|---|---|---|---|---|---|---|---|---|---|---|---|---|
|  |  | **VEP** | **MPRA** | **eCLIP** | **Loc-LR** | **Loc-SR** | **HL** | **MRL** | **Pair** | **TE h** | **TE m** | **Prot** | **GO** |
|  |  | **AUPRC** | **Pearson r** | **AUPRC** | **AUPRC** | **AUPRC** | **Pearson r** | **Pearson r** | **Pearson r** | **Pearson r** | **Pearson r** | **AUPRC** | **AUPRC** |
| Naive-4 | baseline | 0.1949 | 0.6429 | 0.4300 | 0.4268 | 0.5958 | 0.1433 | 0.1432 | 0.5099 | 0.2820 | 0.2536 | 0.2147 | 0.2318 |
| Naive-6 | baseline | 0.1933 | 0.6429 | 0.4356 | 0.4283 | 0.6010 | 0.1970 | 0.1440 | 0.5114 | 0.3073 | 0.2803 | 0.2157 | 0.2352 |
| 3UTRBERT | Global mean | 0.3148 | 0.5947 | 0.3110 | 0.4454 | 0.6324 | 0.4071 | 0.1901 | 0.4984 | 0.5647 | 0.5341 | 0.2701 | 0.2751 |
|  | Region-aware | 0.3055 | 0.6796 | 0.3670 | 0.4791 | 0.6891 | 0.5477 | 0.3417 | 0.5859 | 0.7126 | 0.6891 | 0.2994 | 0.3226 |
|  | Equal thirds | 0.3229 | 0.6728 | 0.3890 | 0.5048 | 0.6936 | 0.5404 | 0.3191 | 0.5066 | 0.6606 | 0.6435 | 0.3096 | 0.3268 |
|  | Nucleotide mean | 0.3071 | 0.6549 | 0.3767 | 0.4982 | 0.7146 | 0.5309 | 0.2760 | 0.5118 | 0.6410 | 0.6260 | 0.3454 | 0.3530 |
| AIDO.RNA 1.6B | Global mean | 0.3156 | 0.6328 | 0.4187 | 0.5462 | 0.7334 | 0.5549 | 0.3586 | 0.5266 | 0.6786 | 0.6587 | 0.3426 | 0.3671 |
|  | Region-aware | 0.3034 | 0.6638 | 0.4048 | 0.5214 | 0.6959 | 0.5770 | 0.4390 | 0.5716 | 0.7526 | 0.7171 | 0.2905 | 0.3242 |
|  | Equal thirds | 0.3006 | 0.6608 | 0.4187 | 0.5297 | 0.7029 | 0.5598 | 0.3959 | 0.4938 | 0.6971 | 0.6676 | 0.2950 | 0.3216 |
|  | Nucleotide mean | 0.3150 | 0.6328 | 0.4184 | 0.5464 | 0.7334 | 0.5546 | 0.3585 | 0.4783 | 0.6781 | 0.6581 | 0.3425 | 0.3671 |
| DNABERT-S | Global mean | 0.3061 | 0.5156 | 0.3710 | 0.5448 | 0.6823 | 0.4664 | 0.2914 | 0.3943 | 0.6342 | 0.5833 | 0.3115 | 0.3177 |
|  | Region-aware | 0.3104 | 0.5705 | 0.3790 | 0.5238 | 0.6533 | 0.4911 | 0.3339 | 0.5084 | 0.6871 | 0.6354 | 0.2781 | 0.3007 |
|  | Equal thirds | 0.3110 | 0.5580 | 0.3789 | 0.5267 | 0.6635 | 0.4746 | 0.3115 | 0.4564 | 0.6336 | 0.5879 | 0.2822 | 0.3000 |
|  | Nucleotide mean | 0.3050 | 0.5166 | 0.3710 | 0.5447 | 0.6818 | 0.4659 | 0.2895 | 0.3803 | 0.6335 | 0.5826 | 0.3120 | 0.3174 |
| DNABERT-2 | Global mean | 0.2615 | 0.5694 | 0.3700 | 0.5470 | 0.6734 | 0.4749 | 0.2850 | 0.5330 | 0.6269 | 0.5865 | 0.3252 | 0.3262 |
|  | Region-aware | **0.3278** | 0.5988 | 0.3903 | 0.5448 | 0.6801 | 0.5039 | 0.3422 | **0.6342** | 0.6948 | 0.6541 | 0.3180 | 0.3303 |
|  | Equal thirds | 0.3102 | 0.5908 | 0.3977 | 0.5463 | 0.6907 | 0.4842 | 0.3219 | 0.5763 | 0.6320 | 0.5973 | 0.3188 | 0.3335 |
|  | Nucleotide mean | 0.2617 | 0.5673 | 0.3697 | 0.5456 | 0.6728 | 0.4709 | 0.2803 | 0.5377 | 0.6236 | 0.5841 | 0.3248 | 0.3255 |
| ERNIE-RNA | Global mean | 0.2905 | 0.5794 | 0.3738 | 0.5297 | 0.6910 | 0.5184 | 0.3280 | 0.5049 | 0.6607 | 0.6314 | 0.3382 | 0.3420 |
|  | Region-aware | 0.3144 | 0.6314 | 0.3977 | 0.5472 | 0.7113 | 0.5452 | 0.3994 | 0.5923 | 0.7298 | 0.6938 | 0.3404 | 0.3551 |
|  | Equal thirds | 0.3207 | 0.6165 | 0.4046 | 0.5499 | 0.7061 | 0.5311 | 0.3779 | 0.5630 | 0.6799 | 0.6496 | 0.3304 | 0.3416 |
|  | Nucleotide mean | 0.2907 | 0.5794 | 0.3737 | 0.5297 | 0.6911 | 0.5183 | 0.3280 | 0.5049 | 0.6603 | 0.6312 | 0.3383 | 0.3416 |
| Evo 1 | Global mean | 0.1219 | 0.2518 | 0.2660 | 0.4515 | 0.6346 | 0.3199 | 0.1874 | 0.2346 | 0.4932 | 0.4497 | 0.2963 | 0.2794 |
|  | Region-aware | 0.1506 | 0.3870 | 0.3060 | 0.4930 | 0.6688 | 0.4351 | 0.2667 | 0.3165 | 0.6042 | 0.5403 | 0.3154 | 0.3153 |
|  | Equal thirds | 0.1464 | 0.3878 | 0.2959 | 0.4718 | 0.6550 | 0.4544 | 0.2700 | 0.3893 | 0.6408 | 0.5928 | 0.3048 | 0.2923 |
|  | Nucleotide mean | 0.1192 | 0.3601 | 0.2644 | 0.4511 | 0.6343 | 0.4366 | 0.2521 | 0.3424 | 0.6158 | 0.5720 | 0.2959 | 0.2790 |
| Evo 2 7B | Global mean | 0.3229 | 0.7048 | **0.4736** | 0.5938 | 0.7671 | 0.6607 | 0.4549 | 0.5055 | 0.7662 | **0.7497** | **0.4079** | **0.4672** |
|  | Region-aware | 0.3020 | 0.7357 | 0.4658 | 0.5498 | 0.7461 | 0.6620 | **0.4584** | 0.6044 | **0.8080** | **0.7858** | 0.3536 | 0.4352 |
|  | Equal thirds | 0.3009 | 0.7372 | **0.4770** | 0.5538 | 0.7356 | 0.6568 | **0.4608** | 0.5507 | **0.7733** | 0.7494 | 0.3506 | 0.4294 |
|  | Nucleotide mean | 0.3225 | 0.7047 | 0.4715 | 0.5911 | 0.7665 | 0.6602 | 0.4555 | 0.5050 | 0.7653 | 0.7459 | **0.4066** | **0.4668** |
| Helix-mRNA | Global mean | 0.1059 | 0.4411 | 0.2611 | 0.4763 | 0.6185 | 0.3734 | 0.1783 | 0.3111 | 0.6213 | 0.5712 | 0.2923 | 0.2645 |
|  | Region-aware | 0.1448 | 0.5946 | 0.2985 | 0.4865 | 0.6417 | 0.4878 | 0.3328 | 0.4893 | 0.7219 | 0.6606 | 0.3108 | 0.3078 |
|  | Equal thirds | 0.0974 | 0.5881 | 0.2959 | 0.5042 | 0.6314 | 0.4685 | 0.2786 | 0.3947 | 0.6787 | 0.6350 | 0.2997 | 0.2817 |
|  | Nucleotide mean | 0.0901 | 0.5444 | 0.2586 | 0.4680 | 0.6230 | 0.4487 | 0.2508 | 0.4668 | 0.6578 | 0.6163 | 0.2908 | 0.2656 |
| HelixNano-mRNA | Global mean | 0.1003 | 0.4697 | 0.2654 | 0.4995 | 0.5977 | 0.3675 | 0.2800 | 0.2420 | 0.5719 | 0.5191 | 0.2828 | 0.2537 |
|  | Region-aware | 0.1473 | 0.6177 | 0.2914 | 0.5076 | 0.6059 | 0.4442 | 0.3522 | 0.4605 | 0.6978 | 0.6488 | 0.2990 | 0.2849 |
|  | Equal thirds | 0.1346 | 0.6148 | 0.2894 | 0.5252 | 0.6277 | 0.4116 | 0.3166 | 0.3998 | 0.6216 | 0.5705 | 0.2947 | 0.2701 |
|  | Nucleotide mean | 0.1002 | 0.5605 | 0.2654 | 0.4994 | 0.5976 | 0.3903 | 0.2984 | 0.4912 | 0.6047 | 0.5538 | 0.2828 | 0.2540 |
| HyenaDNA | Global mean | 0.2400 | 0.4860 | 0.3765 | 0.5539 | 0.6739 | 0.4600 | 0.3066 | 0.4565 | 0.6384 | 0.5861 | 0.3166 | 0.3158 |
|  | Region-aware | 0.2984 | 0.5077 | 0.4026 | 0.5805 | 0.6923 | 0.4869 | 0.3505 | 0.5647 | 0.7221 | 0.6658 | 0.3246 | 0.3352 |
|  | Equal thirds | 0.2950 | 0.5076 | 0.3926 | 0.5595 | 0.6834 | 0.4655 | 0.3162 | 0.4937 | 0.6512 | 0.5962 | 0.3149 | 0.3210 |
|  | Nucleotide mean | 0.2401 | 0.4860 | 0.3765 | 0.5539 | 0.6740 | 0.4598 | 0.3061 | 0.4539 | 0.6385 | 0.5861 | 0.3166 | 0.3158 |
| NT 2.5B | Global mean | 0.3185 | 0.5982 | 0.4049 | 0.5509 | 0.7220 | 0.5703 | 0.3413 | 0.5696 | 0.7062 | 0.6676 | 0.3195 | 0.3598 |
|  | Region-aware | 0.2753 | 0.6425 | 0.3675 | 0.5116 | 0.6923 | 0.5813 | 0.4074 | 0.6109 | 0.7591 | 0.7235 | 0.2697 | 0.3255 |
|  | Equal thirds | 0.2892 | 0.6427 | 0.3861 | 0.5051 | 0.6945 | 0.5781 | 0.3816 | 0.5424 | 0.7206 | 0.6843 | 0.2696 | 0.3246 |
|  | Nucleotide mean | 0.3182 | 0.5984 | 0.4044 | 0.5487 | 0.7218 | 0.5690 | 0.3477 | 0.5657 | 0.7060 | 0.6673 | 0.3190 | 0.3591 |
| Orthrus large 6-track | Global mean | 0.2757 | 0.6242 | 0.4372 | 0.5990 | **0.7794** | **0.6673** | 0.4278 | 0.5289 | 0.7409 | 0.6999 | 0.3926 | 0.4297 |
|  | Region-aware | 0.3059 | 0.6562 | 0.4407 | **0.5995** | 0.7707 | 0.6663 | 0.4388 | 0.5666 | 0.7705 | 0.7261 | 0.3711 | 0.4223 |
|  | Equal thirds | 0.3176 | 0.6559 | 0.4411 | 0.5959 | 0.7659 | 0.6626 | 0.4157 | 0.5343 | 0.7389 | 0.6985 | 0.3652 | 0.4047 |
|  | Nucleotide mean | 0.2754 | 0.6242 | 0.4371 | **0.5992** | **0.7792** | **0.6672** | 0.4277 | 0.5292 | 0.7411 | 0.6999 | 0.3926 | 0.4286 |
| Orthrus base 4-track | Global mean | 0.2712 | 0.6164 | 0.4128 | 0.5834 | 0.7492 | 0.5413 | 0.3620 | 0.5668 | 0.6899 | 0.6519 | 0.3626 | 0.3775 |
|  | Region-aware | 0.3095 | 0.6453 | 0.4157 | 0.5834 | 0.7482 | 0.5489 | 0.4067 | **0.6770** | 0.7452 | 0.6992 | 0.3469 | 0.3797 |
|  | Equal thirds | 0.3164 | 0.6434 | 0.4197 | 0.5859 | 0.7483 | 0.5472 | 0.3743 | 0.5474 | 0.6953 | 0.6534 | 0.3378 | 0.3624 |
|  | Nucleotide mean | 0.2711 | 0.6164 | 0.4127 | 0.5833 | 0.7493 | 0.5413 | 0.3617 | 0.5670 | 0.6899 | 0.6520 | 0.3625 | 0.3772 |
| RNA-FM | Global mean | 0.2566 | 0.4853 | 0.3524 | 0.5095 | 0.6695 | 0.4750 | 0.2945 | 0.5022 | 0.6176 | 0.5808 | 0.3217 | 0.3220 |
|  | Region-aware | 0.3028 | 0.5573 | 0.3971 | 0.5576 | 0.7103 | 0.5239 | 0.3707 | 0.5899 | 0.7146 | 0.6744 | 0.3408 | 0.3512 |
|  | Equal thirds | 0.3063 | 0.5547 | 0.3953 | 0.5514 | 0.6883 | 0.4966 | 0.3558 | 0.5037 | 0.6530 | 0.6110 | 0.3238 | 0.3316 |
|  | Nucleotide mean | 0.2563 | 0.4852 | 0.3516 | 0.5093 | 0.6694 | 0.4741 | 0.2943 | 0.5021 | 0.6165 | 0.5795 | 0.3217 | 0.3216 |
| RNA-MSM | Global mean | 0.1631 | 0.3445 | 0.2825 | 0.4827 | 0.6375 | 0.3397 | 0.1580 | 0.4728 | 0.5138 | 0.4730 | 0.2963 | 0.2755 |
|  | Region-aware | 0.2423 | 0.4099 | 0.3299 | 0.5372 | 0.6720 | 0.4329 | 0.2827 | 0.5348 | 0.6289 | 0.5307 | 0.3099 | 0.3092 |
|  | Equal thirds | 0.2059 | 0.3860 | 0.3154 | 0.5250 | 0.6513 | 0.3905 | 0.2048 | 0.4715 | 0.5487 | 0.4922 | 0.3003 | 0.2855 |
|  | Nucleotide mean | 0.1637 | 0.3589 | 0.2819 | 0.4824 | 0.6375 | 0.3394 | 0.1583 | 0.4732 | 0.5123 | 0.4720 | 0.2963 | 0.2754 |
| RNABERT | Global mean | 0.1924 | 0.2355 | 0.1925 | 0.3888 | 0.5749 | 0.2063 | 0.0570 | 0.3200 | 0.3108 | 0.2651 | 0.2803 | 0.2379 |
|  | Region-aware | 0.2631 | 0.2598 | 0.2222 | 0.4014 | 0.5733 | 0.3714 | 0.1154 | 0.4253 | 0.4814 | 0.3798 | 0.2872 | 0.2523 |
|  | Equal thirds | 0.2211 | 0.2569 | 0.2097 | 0.4064 | 0.5728 | 0.3288 | 0.1104 | 0.3945 | 0.4570 | 0.4041 | 0.2833 | 0.2405 |
|  | Nucleotide mean | 0.1924 | 0.2443 | 0.1927 | 0.3890 | 0.5750 | 0.2388 | 0.0590 | 0.3194 | 0.3542 | 0.3089 | 0.2804 | 0.2381 |
| RNAErnie | Global mean | 0.2891 | 0.4864 | 0.3441 | 0.4897 | 0.6593 | 0.3986 | 0.1957 | 0.4937 | 0.5620 | 0.5066 | 0.3107 | 0.3060 |
|  | Region-aware | 0.2985 | 0.5209 | 0.3335 | 0.4679 | 0.6525 | 0.4680 | 0.2824 | 0.5568 | 0.6431 | 0.5933 | 0.2725 | 0.2881 |
|  | Equal thirds | 0.3121 | 0.5169 | 0.3473 | 0.4883 | 0.6414 | 0.4284 | 0.2542 | 0.4984 | 0.5965 | 0.5360 | 0.2808 | 0.2871 |
|  | Nucleotide mean | 0.2886 | 0.4864 | 0.3427 | 0.4890 | 0.6592 | 0.3984 | 0.1945 | 0.4940 | 0.5603 | 0.5037 | 0.3108 | 0.3059 |
| RiNALMo giga | Global mean | 0.3153 | 0.7418 | 0.3927 | 0.5919 | 0.6898 | 0.5318 | 0.4169 | 0.4646 | 0.6672 | 0.6357 | 0.3281 | 0.3477 |
|  | Region-aware | 0.3001 | **0.7687** | 0.3839 | 0.5482 | 0.6654 | 0.5442 | 0.4541 | 0.5686 | 0.7316 | 0.6780 | 0.2791 | 0.3076 |
|  | Equal thirds | 0.2921 | **0.7677** | 0.3824 | 0.5460 | 0.6629 | 0.5364 | 0.4249 | 0.5208 | 0.6731 | 0.6353 | 0.2862 | 0.3112 |
|  | Nucleotide mean | 0.3153 | 0.7418 | 0.3926 | 0.5918 | 0.6898 | 0.5317 | 0.4169 | 0.4581 | 0.6672 | 0.6357 | 0.3281 | 0.3475 |
| SpliceBERT | Global mean | 0.2835 | 0.5326 | 0.3771 | 0.5325 | 0.7264 | 0.5309 | 0.3361 | 0.5132 | 0.6699 | 0.6429 | 0.3605 | 0.3822 |
|  | Region-aware | 0.3184 | 0.5901 | 0.4030 | 0.5659 | 0.7464 | 0.5613 | 0.4065 | 0.5542 | 0.7426 | 0.7044 | 0.3614 | 0.3934 |
|  | Equal thirds | **0.3253** | 0.5968 | 0.4091 | 0.5701 | 0.7388 | 0.5468 | 0.3683 | 0.5283 | 0.7000 | 0.6685 | 0.3525 | 0.3834 |
|  | Nucleotide mean | 0.2833 | 0.5324 | 0.3770 | 0.5327 | 0.7264 | 0.5309 | 0.3360 | 0.5132 | 0.6696 | 0.6427 | 0.3604 | 0.3821 |
| UTR-LM | Global mean | 0.2260 | 0.5083 | 0.3320 | 0.5055 | 0.6549 | 0.4342 | 0.2329 | 0.4431 | 0.5696 | 0.5326 | 0.3180 | 0.3117 |
|  | Region-aware | 0.3064 | 0.5659 | 0.3706 | 0.5489 | 0.7007 | 0.5023 | 0.3460 | 0.5014 | 0.6887 | 0.6412 | 0.3359 | 0.3418 |
|  | Equal thirds | 0.2780 | 0.5624 | 0.3751 | 0.5473 | 0.6789 | 0.4761 | 0.2878 | 0.4695 | 0.6240 | 0.5753 | 0.3215 | 0.3213 |
|  | Nucleotide mean | 0.2255 | 0.5083 | 0.3319 | 0.5053 | 0.6549 | 0.4339 | 0.2331 | 0.4455 | 0.5693 | 0.5318 | 0.3181 | 0.3117 |

**Table S4:**
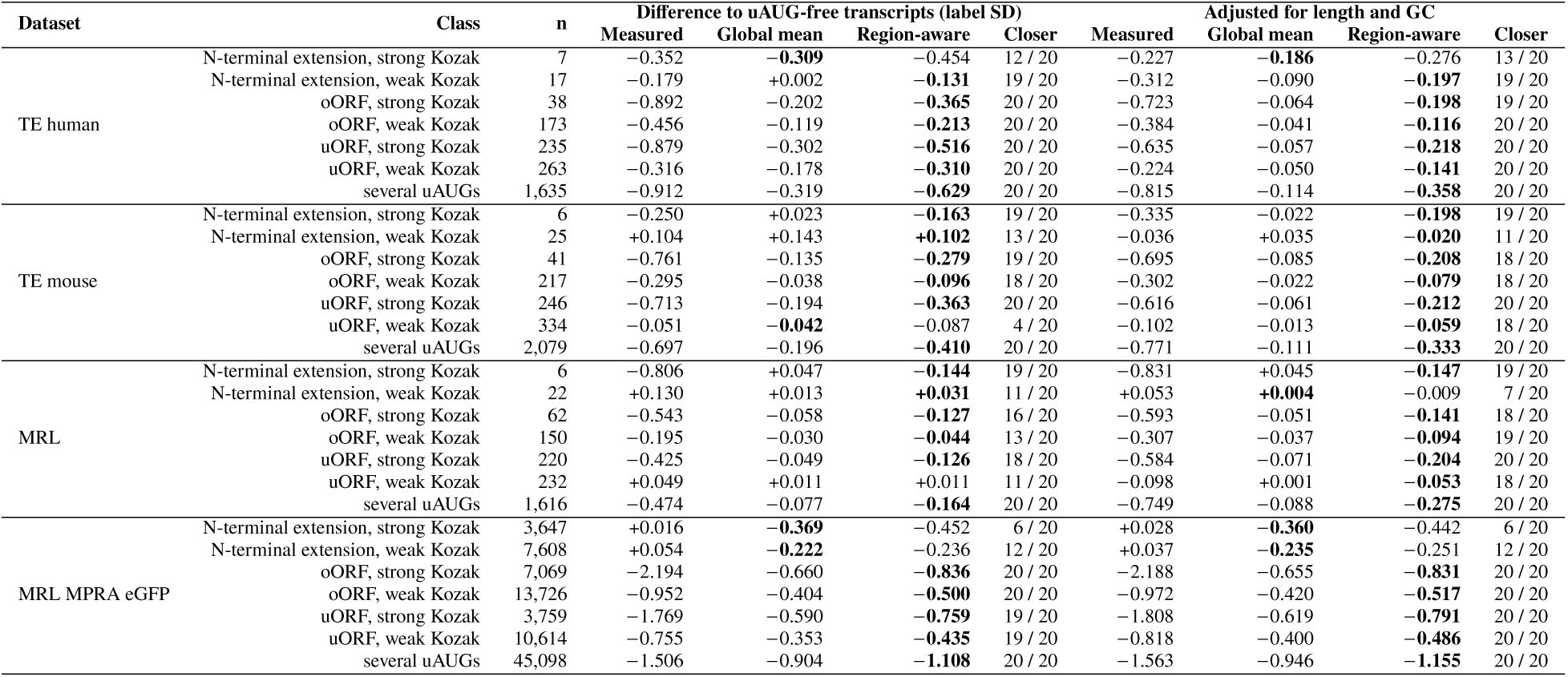
uAUG classes in natural transcripts: measured and predicted differences to uAUG-free transcripts (mean over 20 models), and the number of models in which the region readout is closer. The prediction closer to the measured difference is **bold**.

**Table S5:** uAUG insertion: mean change of the prediction for each uAUG kind and Kozak context, for the control edits and for removal, with each probe’s *ρ* and slope against the dataset’s measured effects (mean over the 20 models of Table S6). In each dataset the prediction closer to the measured effect, the higher *ρ* and the slope closer to 1 are **bold**.

| Dataset | Readout | N-terminal extension |  | oORF |  | uORF |  | Control edit | Ablation | Against measured |  | 5′ UTR share | Closer |
|---|---|---|---|---|---|---|---|---|---|---|---|---|---|
|  |  | strong Kozak | weak Kozak | strong Kozak | weak Kozak | strong Kozak | weak Kozak |  |  | *ρ* | Slope |  |  |
| TE human | Measured (n) | −0.35 (7) | −0.09 (26) | −0.94 (57) | −0.47 (242) | −0.90 (348) | −0.33 (387) |  |  |  |  |  |  |
|  | Global mean | −0.000 | +0.004 | −0.003 | +0.005 | −0.003 | +0.007 | +0.007 | +0.004 | +0.35 | +0.00 |  |  |
|  | Region-aware | **−0.099** | **−0.034** | **−0.105** | **−0.030** | **−0.104** | **−0.027** | +0.013 | +0.058 | **+0.51** | **+0.12** | 0.56 | 18 / 20 |
| MRL | Measured (n) | −0.52 (7) | −0.03 (28) | −0.63 (90) | −0.21 (256) | −0.48 (342) | −0.05 (403) |  |  |  |  |  |  |
|  | Global mean | −0.014 | −0.006 | −0.016 | −0.006 | −0.015 | −0.006 | +0.002 | +0.012 | +0.49 | +0.03 |  |  |
|  | Region-aware | **−0.100** | **−0.037** | **−0.102** | **−0.035** | **−0.099** | **−0.035** | +0.008 | +0.059 | **+0.68** | **+0.18** | 1.00 | 18 / 20 |
| MRL MPRA eGFP | Measured (n) | +0.01 (4,506) | +0.05 (9,443) | −2.20 (8,875) | −0.95 (17,018) | −1.77 (4,691) | −0.76 (13,199) |  |  |  |  |  |  |
|  | Global mean | **−0.341** | **−0.256** | −0.668 | −0.373 | −0.623 | −0.350 | −0.023 | +0.454 | +0.84 | +0.34 |  |  |
|  | Region-aware | −0.425 | −0.273 | **−0.848** | **−0.457** | **−0.799** | **−0.415** | −0.014 | +0.560 | **+0.88** | **+0.42** | 0.50 | 20 / 20 |

**Table S6:**
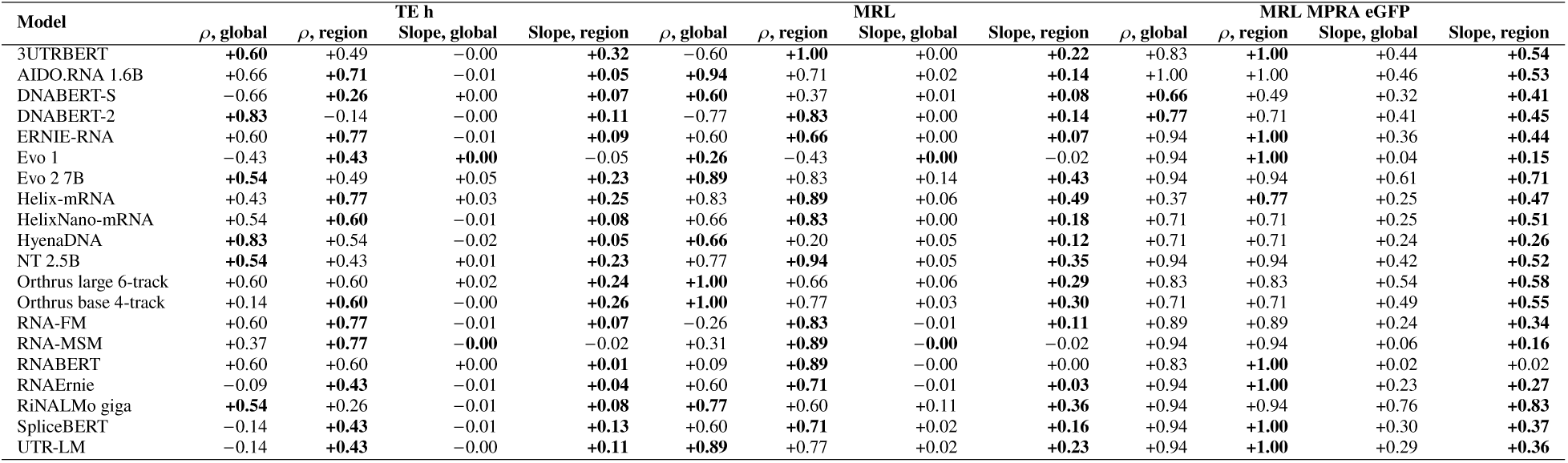
uAUG insertion per model: Spearman *ρ* and slope of the predicted change against the dataset’s measured class effects. In each pair the higher *ρ* and the slope closer to 1 are **bold**. TE h, TE human.

**Table S7:** HL edits: mean change of the predicted half-life for codon recoding (a quarter or all of the codons) and for 3^′^ UTR insertions.

| Model | Readout | 25% of the codons |  |  | All codons |  |  | 3′ UTR, four copies |  |  |  | CDS share |
|---|---|---|---|---|---|---|---|---|---|---|---|---|
|  |  | Optimal | Non-optimal | Random | Optimal | Non-optimal | Random | ARE | Shuffled ARE | let-7 8mer | Shuffled let-7 | (non-optimal, 25%) |
| 3UTRBERT | Global mean | +0.084 | −0.147 | −0.041 | +0.435 | −0.121 | −0.124 | −0.047 | −0.035 | +0.011 | +0.019 |  |
|  | Region-aware | +0.100 | −0.055 | −0.010 | +0.405 | +0.153 | −0.002 | −0.202 | −0.134 | +0.071 | +0.035 | 0.31 |
| AIDO.RNA 1.6B | Global mean | +0.137 | −0.313 | −0.081 | +0.580 | −0.874 | −0.536 | −0.068 | −0.067 | +0.033 | +0.034 |  |
|  | Region-aware | +0.104 | −0.261 | −0.083 | +0.611 | −0.348 | −0.378 | −0.095 | −0.074 | +0.065 | +0.061 | 0.87 |
| DNABERT-S | Global mean | +0.108 | −0.077 | −0.034 | +0.356 | +0.073 | −0.073 | −0.031 | −0.037 | +0.003 | +0.011 |  |
|  | Region-aware | +0.067 | −0.004 | +0.009 | +0.230 | −0.088 | +0.048 | −0.057 | −0.063 | +0.017 | +0.024 | -11.93 |
| DNABERT-2 | Global mean | +0.108 | −0.098 | −0.032 | +0.682 | −0.156 | −0.159 | −0.037 | −0.030 | −0.008 | −0.000 |  |
|  | Region-aware | +0.045 | +0.028 | +0.005 | +0.341 | +0.198 | −0.002 | −0.098 | −0.067 | +0.032 | +0.040 | 2.27 |
| ERNIE-RNA | Global mean | +0.209 | −0.204 | −0.033 | +0.545 | −1.264 | −0.620 | −0.105 | −0.094 | +0.035 | +0.018 |  |
|  | Region-aware | +0.113 | −0.078 | −0.015 | +0.229 | −0.564 | −0.332 | −0.136 | −0.116 | +0.047 | +0.052 | 0.85 |
| Evo 1 | Global mean | +0.117 | −0.132 | −0.181 | +1.357 | −0.292 | −0.114 | +0.054 | +0.075 | +0.077 | +0.152 |  |
|  | Region-aware | +0.136 | +0.140 | +0.091 | +0.765 | +0.266 | +0.283 | −0.087 | −0.098 | +0.052 | +0.043 | 0.80 |
| Evo 2 7B | Global mean | +0.036 | −0.220 | −0.051 | +0.454 | +0.116 | −0.288 | −0.136 | −0.054 | +0.017 | +0.034 |  |
|  | Region-aware | +0.018 | −0.182 | −0.047 | +0.516 | +0.026 | −0.129 | −0.158 | −0.061 | +0.018 | +0.036 | 0.51 |
| Helix-mRNA | Global mean | −0.029 | +0.040 | +0.007 | −0.015 | +0.153 | +0.053 | −0.005 | −0.014 | +0.012 | +0.013 |  |
|  | Region-aware | +0.057 | +0.024 | +0.032 | +0.127 | −0.000 | +0.067 | −0.122 | −0.098 | +0.054 | +0.050 | 3.48 |
| HelixNano-mRNA | Global mean | +0.041 | +0.007 | +0.005 | +0.190 | −0.047 | +0.005 | +0.008 | +0.005 | −0.003 | −0.004 |  |
|  | Region-aware | +0.062 | +0.080 | +0.038 | +0.234 | +0.077 | +0.104 | −0.025 | −0.028 | +0.039 | +0.040 | 1.01 |
| HyenaDNA | Global mean | +0.158 | −0.048 | +0.007 | +0.646 | −0.374 | −0.063 | −0.077 | −0.092 | +0.014 | +0.020 |  |
|  | Region-aware | +0.123 | +0.025 | +0.020 | +0.537 | −0.024 | −0.019 | −0.080 | −0.086 | +0.051 | +0.062 | 1.90 |
| NT 2.5B | Global mean | +0.131 | −0.128 | −0.019 | +0.658 | −0.440 | −0.087 | −0.091 | −0.058 | +0.034 | +0.031 |  |
|  | Region-aware | +0.100 | −0.039 | −0.011 | +0.598 | −0.116 | −0.045 | −0.033 | −0.050 | +0.053 | +0.046 | 0.66 |
| Orthrus large 6-track | Global mean | +0.146 | −0.234 | −0.048 | +0.968 | −0.792 | −0.232 | −0.003 | −0.002 | +0.004 | +0.004 |  |
|  | Region-aware | +0.149 | −0.208 | −0.039 | +1.005 | −0.629 | −0.130 | −0.009 | −0.019 | +0.011 | +0.012 | 0.78 |
| Orthrus base 4-track | Global mean | +0.077 | −0.204 | −0.081 | +0.385 | −0.234 | −0.411 | −0.009 | −0.016 | +0.019 | +0.024 |  |
|  | Region-aware | +0.074 | −0.156 | −0.054 | +0.331 | −0.116 | −0.265 | −0.039 | −0.045 | +0.037 | +0.037 | 0.97 |
| RNA-FM | Global mean | +0.139 | −0.156 | −0.044 | +0.408 | −0.548 | −0.232 | −0.058 | −0.053 | +0.020 | +0.023 |  |
|  | Region-aware | +0.053 | −0.056 | −0.026 | +0.088 | −0.251 | −0.227 | −0.069 | −0.068 | +0.073 | +0.061 | 0.05 |
| RNA-MSM | Global mean | +0.056 | +0.017 | +0.011 | +0.143 | +0.079 | +0.004 | −0.014 | −0.015 | +0.004 | +0.004 |  |
|  | Region-aware | +0.044 | +0.159 | +0.042 | +0.105 | +0.667 | +0.123 | −0.057 | −0.057 | +0.060 | +0.061 | 1.21 |
| RNABERT | Global mean | +0.049 | −0.002 | +0.006 | +0.164 | −0.104 | +0.017 | −0.004 | −0.003 | −0.011 | −0.009 |  |
|  | Region-aware | +0.044 | +0.152 | +0.042 | +0.112 | +0.489 | +0.155 | −0.049 | −0.048 | +0.062 | +0.060 | 1.18 |
| RNAErnie | Global mean | +0.131 | −0.097 | −0.017 | +0.570 | −0.390 | −0.070 | −0.063 | −0.075 | +0.029 | +0.027 |  |
|  | Region-aware | +0.075 | +0.132 | +0.034 | +0.245 | +0.655 | +0.133 | −0.066 | −0.071 | +0.076 | +0.074 | 1.15 |
| RiNALMo giga | Global mean | +0.049 | −0.170 | −0.054 | +0.268 | −0.972 | −0.362 | −0.075 | −0.066 | +0.020 | +0.029 |  |
|  | Region-aware | +0.057 | −0.121 | −0.054 | +0.365 | −0.452 | −0.325 | −0.099 | −0.089 | +0.030 | +0.042 | 0.35 |
| SpliceBERT | Global mean | +0.132 | −0.306 | −0.095 | +0.514 | −1.309 | −0.675 | −0.113 | −0.086 | +0.040 | +0.022 |  |
|  | Region-aware | +0.059 | −0.144 | −0.064 | +0.229 | −0.347 | −0.402 | −0.132 | −0.108 | +0.051 | +0.047 | 0.50 |
| UTR-LM | Global mean | +0.113 | −0.030 | −0.008 | +0.251 | +0.110 | −0.103 | −0.067 | −0.055 | +0.004 | +0.013 |  |
|  | Region-aware | +0.072 | +0.023 | +0.010 | +0.186 | −0.059 | −0.103 | −0.101 | −0.089 | +0.059 | +0.053 | 1.62 |

**Table S8:** Natural element groupings: measured and predicted differences (mean over 20 models), raw and adjusted, with per-model verdicts. The prediction closer to the measured difference is **bold**.

| Element | Dataset | Expected | n (with / without) | Difference |  |  |  | Adjusted for length and GC |  |  |  |
|---|---|---|---|---|---|---|---|---|---|---|---|
|  |  |  |  | Measured | Global mean | Region-aware | Region / global / tie | Measured | Global mean | Region-aware | Region / global / tie |
| out-of-frame uAUG vs none | TE human | lower | 3,962 / 5,699 | −0.817 | −0.290 | **−0.537** | 20 / 0 / 0 | −0.653 | −0.088 | **−0.258** | 20 / 0 / 0 |
|  | TE mouse | lower | 4,906 / 5,147 | −0.573 | −0.159 | **−0.334** | 20 / 0 / 0 | −0.528 | −0.072 | **−0.219** | 20 / 0 / 0 |
|  | MRL | lower | 4,252 / 5,039 | −0.412 | −0.061 | **−0.126** | 19 / 0 / 1 | −0.603 | −0.068 | **−0.200** | 20 / 0 / 0 |
|  | MPRA eGFP | lower | 125,264 / 143,310 | −1.544 | −0.670 | **−0.835** | 20 / 0 / 0 | −1.584 | −0.696 | **−0.864** | 20 / 0 / 0 |
|  | MPRA mCherry | lower | 74,050 / 86,387 | −1.368 | −0.574 | **−0.720** | 20 / 0 / 0 | −1.399 | −0.592 | **−0.740** | 20 / 0 / 0 |
|  | MPRA designed | lower | 22,562 / 65,242 | −1.475 | −0.608 | **−0.784** | 20 / 0 / 0 | −1.511 | −0.643 | **−0.819** | 20 / 0 / 0 |
|  | MPRA varying | lower | 42,083 / 51,704 | −1.364 | −0.563 | **−0.706** | 20 / 0 / 0 | −1.502 | −0.607 | **−0.768** | 20 / 0 / 0 |
| in-frame uAUG vs none | TE human | – | 616 / 5,699 | −0.579 | −0.271 | **−0.468** | 20 / 0 / 0 | −0.387 | −0.056 | **−0.209** | 20 / 0 / 0 |
|  | TE mouse | – | 800 / 5,147 | −0.345 | −0.104 | **−0.236** | 20 / 0 / 0 | −0.368 | −0.049 | **−0.192** | 20 / 0 / 0 |
|  | MRL | – | 641 / 5,039 | −0.201 | −0.017 | **−0.060** | 10 / 1 / 9 | −0.364 | −0.035 | **−0.147** | 20 / 0 / 0 |
|  | MPRA eGFP | – | 49,894 / 143,310 | −0.391 | **−0.363** | −0.430 | 8 / 12 / 0 | −0.415 | **−0.383** | −0.451 | 8 / 12 / 0 |
|  | MPRA mCherry | – | 31,360 / 86,387 | −0.333 | −0.301 | **−0.358** | 8 / 11 / 1 | −0.348 | −0.313 | **−0.371** | 8 / 11 / 1 |
|  | MPRA designed | – | 12,213 / 65,242 | −0.056 | **−0.046** | −0.092 | 5 / 10 / 5 | −0.332 | **−0.321** | −0.371 | 5 / 10 / 5 |
|  | MPRA varying | – | 12,743 / 51,704 | −0.228 | **−0.311** | −0.384 | 3 / 16 / 1 | −0.353 | **−0.371** | −0.456 | 3 / 13 / 4 |
| CAI, top vs bottom quarter | TE human | higher | 2,570 / 2,570 | −0.138 | −0.241 | **−0.200** | 11 / 0 / 9 | +0.184 | +0.103 | **+0.107** | 2 / 0 / 18 |
|  | TE mouse | higher | 2,714 / 2,714 | +0.145 | −0.057 | **+0.018** | 13 / 1 / 6 | +0.333 | +0.116 | **+0.188** | 13 / 0 / 7 |
|  | MRL | higher | 2,484 / 2,483 | +0.095 | **+0.061** | +0.049 | 0 / 1 / 19 | −0.030 | **+0.024** | +0.028 | 0 / 0 / 20 |
|  | HL human | higher | 3,235 / 3,235 | +0.235 | +0.256 | **+0.218** | 1 / 0 / 19 | +0.184 | **+0.188** | +0.135 | 0 / 1 / 19 |
|  | HL mouse | higher | 3,431 / 3,431 | +0.269 | **+0.179** | +0.171 | 3 / 9 / 8 | +0.276 | **+0.183** | +0.166 | 4 / 4 / 12 |
| codon optimality (CSC), top vs bottom quarter | TE human | higher | 2,570 / 2,570 | +0.425 | +0.157 | **+0.257** | 17 / 0 / 3 | +0.541 | +0.345 | **+0.427** | 9 / 1 / 10 |
|  | TE mouse | higher | 2,714 / 2,714 | +0.704 | +0.385 | **+0.502** | 17 / 0 / 3 | +0.561 | +0.333 | **+0.412** | 10 / 0 / 10 |
|  | MRL | higher | 2,483 / 2,483 | +0.231 | +0.170 | **+0.178** | 5 / 0 / 15 | +0.243 | +0.181 | **+0.241** | 3 / 0 / 17 |
|  | HL human | higher | 3,235 / 3,235 | +0.787 | +0.552 | **+0.585** | 10 / 4 / 6 | +0.929 | +0.575 | **+0.650** | 9 / 0 / 11 |
|  | HL mouse | higher | 3,431 / 3,431 | +0.860 | +0.546 | **+0.628** | 15 / 3 / 2 | +0.827 | +0.489 | **+0.577** | 11 / 1 / 8 |
| AUUUA, ≥3 vs none | HL human | lower | 5,179 / 4,216 | −0.531 | −0.443 | **−0.463** | 10 / 3 / 7 | −0.263 | **−0.179** | −0.161 | 2 / 5 / 13 |
|  | HL mouse | lower | 4,370 / 5,034 | −0.558 | −0.425 | **−0.445** | 8 / 3 / 9 | −0.227 | **−0.113** | −0.073 | 0 / 17 / 3 |
| let-7 site vs none | HL human | lower | 1,982 / 10,957 | −0.255 | **−0.178** | −0.162 | 1 / 9 / 10 | −0.057 | **+0.007** | +0.028 | 0 / 11 / 9 |
|  | HL mouse | lower | 1,913 / 11,810 | −0.227 | **−0.159** | −0.151 | 2 / 9 / 9 | −0.008 | **+0.025** | +0.046 | 0 / 8 / 12 |
| miR-16 site vs none | HL human | lower | 3,090 / 9,849 | −0.208 | **−0.161** | −0.146 | 1 / 4 / 15 | −0.045 | **−0.006** | +0.007 | 1 / 1 / 18 |
|  | HL mouse | lower | 3,179 / 10,544 | −0.301 | −0.193 | **−0.203** | 8 / 3 / 9 | −0.101 | −0.011 | **−0.016** | 5 / 1 / 14 |

**Table S9:**
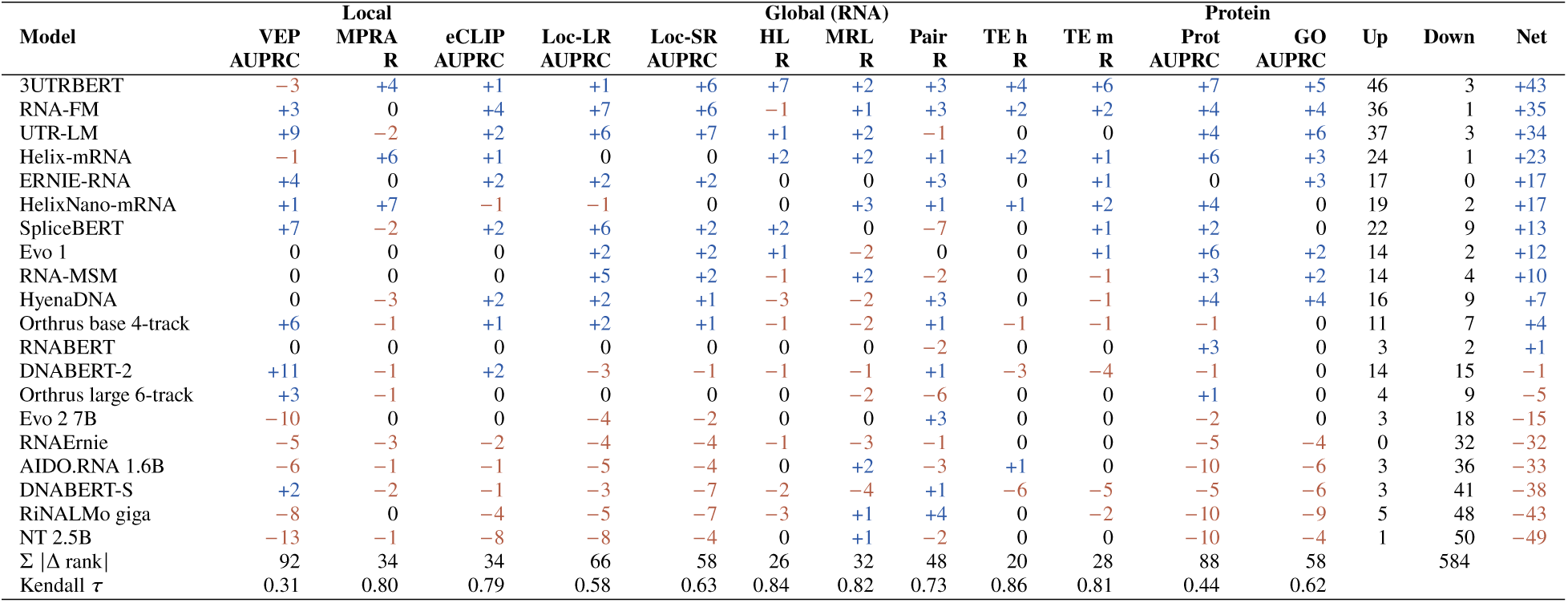
Rank of each model under the global mean minus its rank under the region-aware readout, per column (positive: moved up, negative: moved down); places moved up and down and the net change over the twelve columns; Σ |Δ rank| per column and Kendall *τ* between the two orders. Short column names as in Table 5.

|  | **Local** |  |  | **Global (RNA)** |  |  |  |  |  |  | **Protein** |  |  |  |  |
|---|---|---|---|---|---|---|---|---|---|---|---|---|---|---|---|
| **Model** | **VEP** | **MPRA** | **eCLIP** | **Loc-LR** | **Loc-SR** | **HL** | **MRL** | **Pair** | **TE h** | **TE m** | **Prot** | **GO** | **Up** | **Down** | **Net** |
|  | **AUPRC** | **R** | **AUPRC** | **AUPRC** | **AUPRC** | **R** | **R** | **R** | **R** | **R** | **AUPRC** | **AUPRC** |  |  |  |
| 3UTRBERT | −3 | +4 | +1 | +1 | +6 | +7 | +2 | +3 | +4 | +6 | +7 | +5 | 46 | 3 | +43 |
| RNA-FM | +3 | 0 | +4 | +7 | +6 | −1 | +1 | +3 | +2 | +2 | +4 | +4 | 36 | 1 | +35 |
| UTR-LM | +9 | −2 | +2 | +6 | +7 | +1 | +2 | −1 | 0 | 0 | +4 | +6 | 37 | 3 | +34 |
| Helix-mRNA | −1 | +6 | +1 | 0 | 0 | +2 | +2 | +1 | +2 | +1 | +6 | +3 | 24 | 1 | +23 |
| ERNIE-RNA | +4 | 0 | +2 | +2 | +2 | 0 | 0 | +3 | 0 | +1 | 0 | +3 | 17 | 0 | +17 |
| HelixNano-mRNA | +1 | +7 | −1 | −1 | 0 | 0 | +3 | +1 | +1 | +2 | +4 | 0 | 19 | 2 | +17 |
| SpliceBERT | +7 | −2 | +2 | +6 | +2 | +2 | 0 | −7 | 0 | +1 | +2 | 0 | 22 | 9 | +13 |
| Evo 1 | 0 | 0 | 0 | +2 | +2 | +1 | −2 | 0 | 0 | +1 | +6 | +2 | 14 | 2 | +12 |
| RNA-MSM | 0 | 0 | 0 | +5 | +2 | −1 | +2 | −2 | 0 | −1 | +3 | +2 | 14 | 4 | +10 |
| HyenaDNA | 0 | −3 | +2 | +2 | +1 | −3 | −2 | +3 | 0 | −1 | +4 | +4 | 16 | 9 | +7 |
| Orthrus base 4-track | +6 | −1 | +1 | +2 | +1 | −1 | −2 | +1 | −1 | −1 | −1 | 0 | 11 | 7 | +4 |
| RNABERT | 0 | 0 | 0 | 0 | 0 | 0 | 0 | −2 | 0 | 0 | +3 | 0 | 3 | 2 | +1 |
| DNABERT-2 | +11 | −1 | +2 | −3 | −1 | −1 | −1 | +1 | −3 | −4 | −1 | 0 | 14 | 15 | −1 |
| Orthrus large 6-track | +3 | −1 | 0 | 0 | 0 | 0 | −2 | −6 | 0 | 0 | +1 | 0 | 4 | 9 | −5 |
| Evo 2 7B | −10 | 0 | 0 | −4 | −2 | 0 | 0 | +3 | 0 | 0 | −2 | 0 | 3 | 18 | −15 |
| RNAErnie | −5 | −3 | −2 | −4 | −4 | −1 | −3 | −1 | 0 | 0 | −5 | −4 | 0 | 32 | −32 |
| AIDO.RNA 1.6B | −6 | −1 | −1 | −5 | −4 | 0 | +2 | −3 | +1 | 0 | −10 | −6 | 3 | 36 | −33 |
| DNABERT-S | +2 | −2 | −1 | −3 | −7 | −2 | −4 | +1 | −6 | −5 | −5 | −6 | 3 | 41 | −38 |
| RiNALMo giga | −8 | 0 | −4 | −5 | −7 | −3 | +1 | +4 | 0 | −2 | −10 | −9 | 5 | 48 | −43 |
| NT 2.5B | −13 | −1 | −8 | −8 | −4 | 0 | +1 | −2 | 0 | 0 | −10 | −4 | 1 | 50 | −49 |
| Σ \|Δ rank\| | 92 | 34 | 34 | 66 | 58 | 26 | 32 | 48 | 20 | 28 | 88 | 58 |  | 584 |  |
| Kendall *τ* | 0.31 | 0.80 | 0.79 | 0.58 | 0.63 | 0.84 | 0.82 | 0.73 | 0.86 | 0.81 | 0.44 | 0.62 |  |  |  |

**Figure S1:**
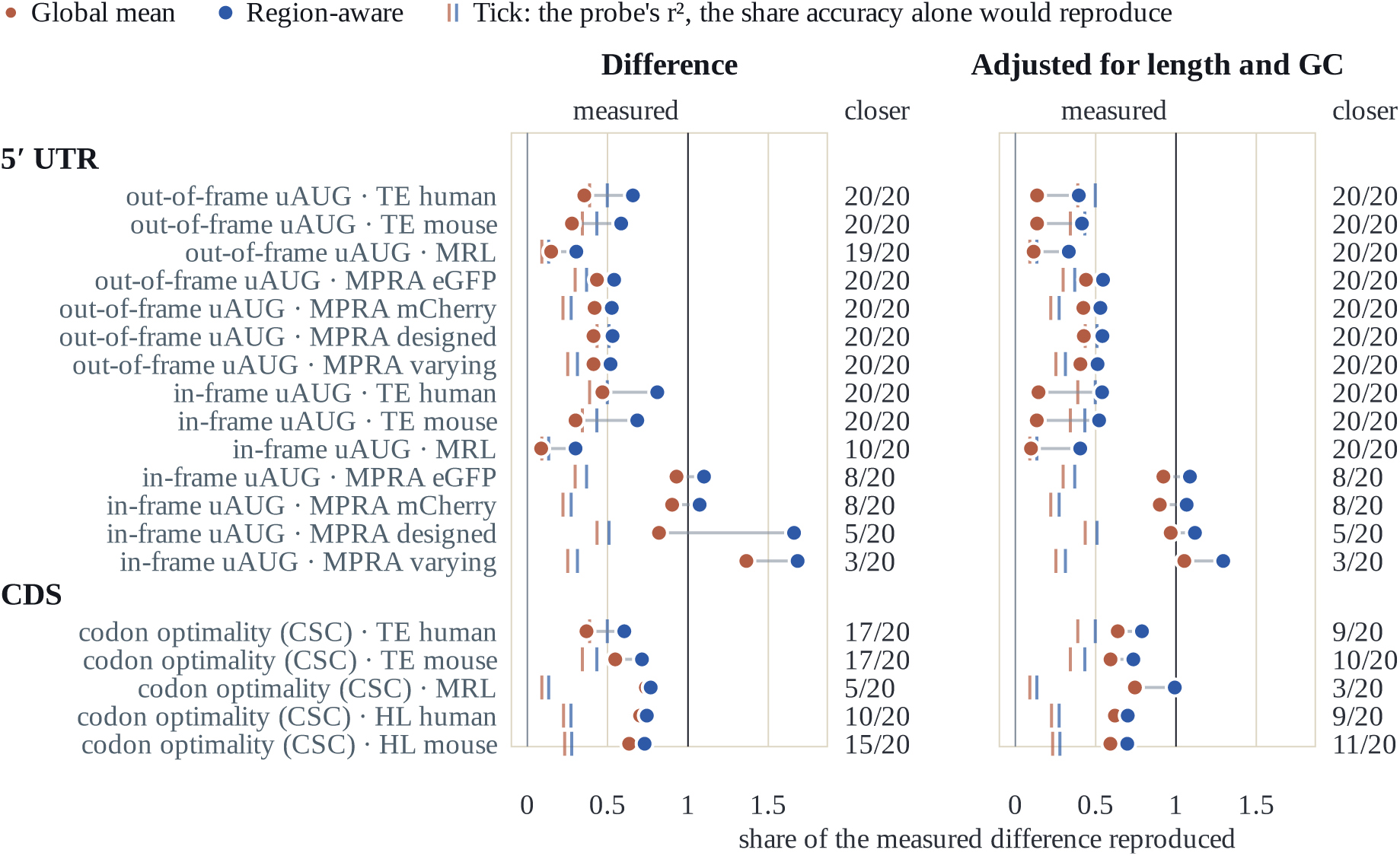
Upstream AUGs and codon optimality in natural transcripts, displayed as in Figure 4. Each row compares two groups of test transcripts: those carrying an out-of-frame or an in-frame uAUG against the uAUG-free ones, and the top against the bottom quartile of the mean codon stability coefficient (CSC) of the CDS. Here the right-hand column counts the models in which the region readout’s error on the group difference is the smaller at Wilcoxon p < 0.05 over the ten splits. Table S8 lists every grouping, the AU-rich elements and microRNA sites included.

**Figure S2:**
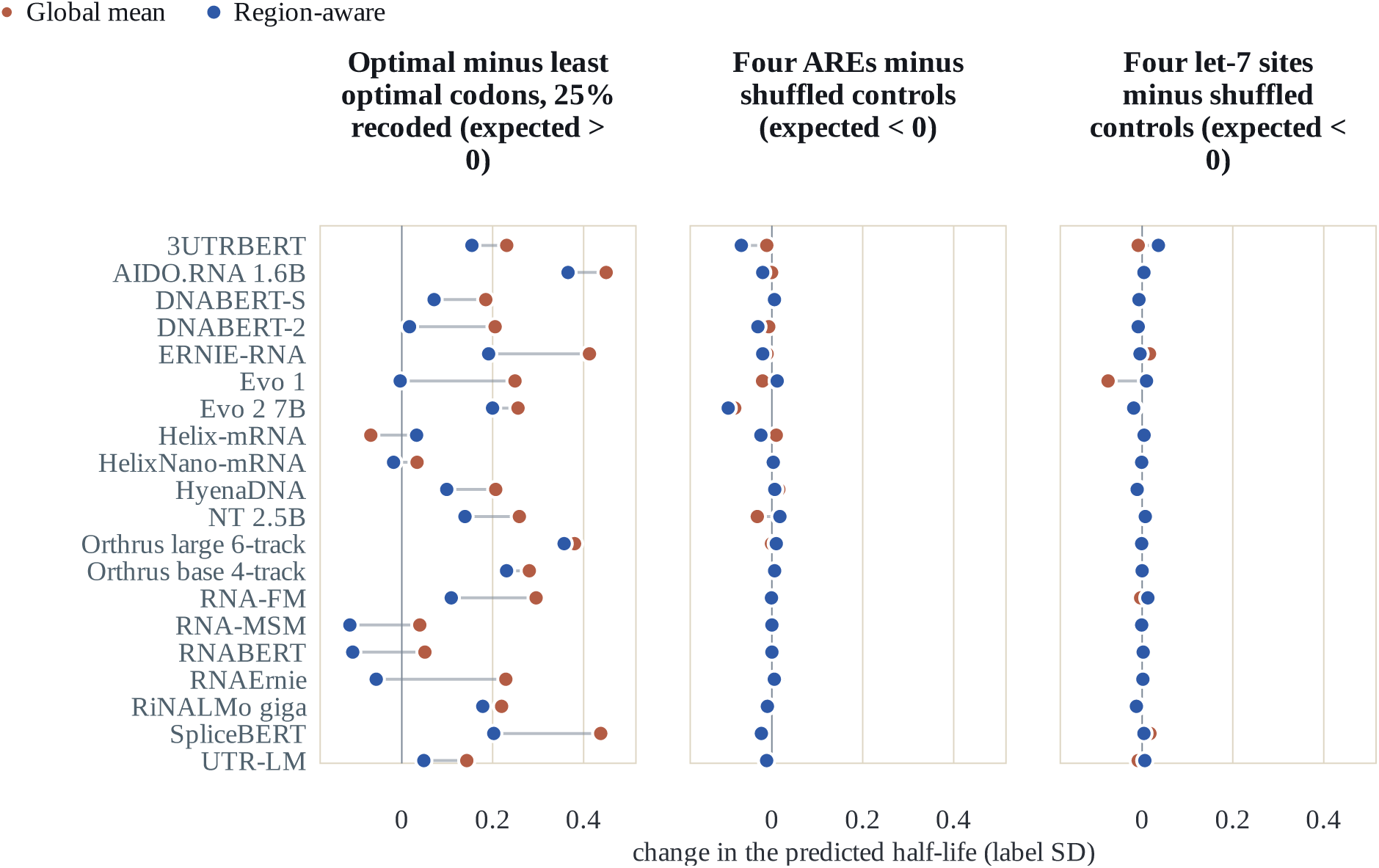
Contrasts of the HL edits for each model: optimal minus least optimal synonyms with a quarter of the codons recoded, and four AREs or let-7 sites minus their shuffled controls, in units of the label’s SD.

**Figure S3:**
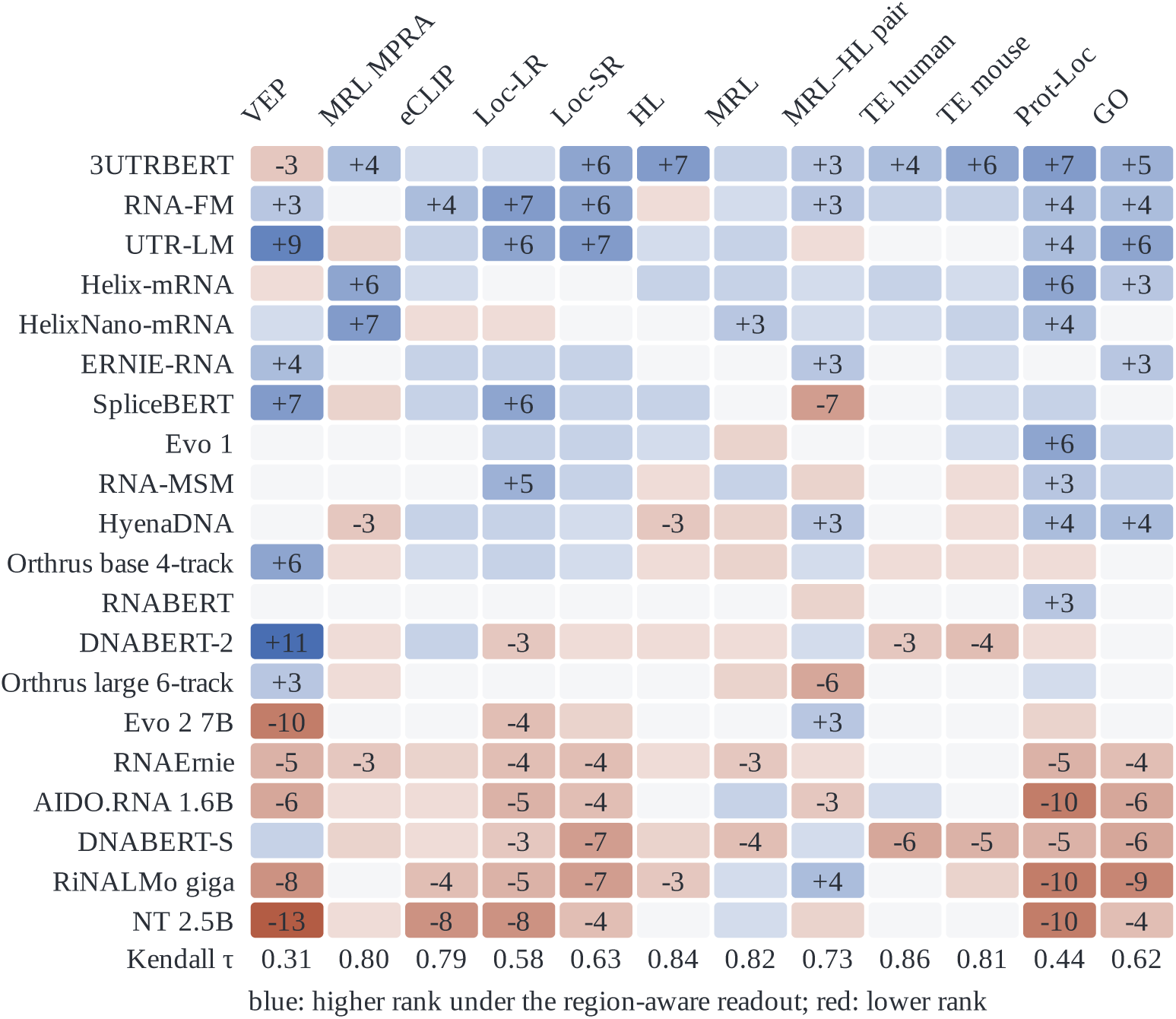
Change in model rank between the global mean and region-aware readouts. Each cell is a model’s rank under the global mean minus its rank under the region-aware readout, within one column over the 20 models: blue, the region readout places it higher; red, lower. Changes of three places or more carry the number. The last row is Kendall *τ* between the two orderings.

